# Cryptic Evolution of Heteroresistance as Adaptation to Treatment Interruptions

**DOI:** 10.64898/2026.05.18.725909

**Authors:** Muqing Ma, Minchae Kang, Tai Do, Minsu Kim

## Abstract

The evolution of antibiotic resistance is traditionally understood as a selective sweep to fixation, yielding easily detectable, population-wide resistance. Many clinical isolates, however, exhibit a subtle phenotype in which resistance remains hidden within a susceptible majority despite a clonal genetic background: a phenomenon clinically recognized as heteroresistance (HR). Treatment failure driven by HR has been widely reported across bacterial and fungal infections and in cancer therapy. To understand when and how HR evolves, and why it is selected over classical population-wide resistance, we conducted de novo evolution experiments starting from susceptible *Escherichia coli* and analyzed the genetic changes and fitness effects in the evolved strains. Prolonged gaps in antibiotic exposure are required for HR to evolve, implicating treatment interruptions as a key driver. HR emerges rapidly and reproducibly with minimal antibiotic use, yet its emergence is not readily detected by routine susceptibility testing. Unlike classical resistance, an evolved HR population partitions at the single-cell level into multiple phenotypes with distinct growth–resistance trade-offs. Their relative abundance shifts dynamically with antibiotic exposure, enabling robust population survival while avoiding the constitutive fitness burden associated with classical resistance. Despite this phenotypic flexibility, stable single mutations including a missense substitution and a short in-frame deletion are sufficient to generate HR, indicating a low evolutionary barrier. Additionally, we found that clinical isolates exhibit genetic and fitness signatures resembling those of our lab-evolved strains, suggesting that clinical HR emerges through the selective mechanism uncovered in our experiments. Together, our results establish HR as a readily evolvable adaptive strategy under treatment interruptions that leverages phenotypic flexibility to maintain resistance at minimal fitness cost, providing mechanistic insight into its emergence and prevalence.

## Introduction

Antibiotic resistance is one of the most urgent threats to global health, undermining the efficacy of life-saving treatments. Infections that were once easily cured are becoming increasingly difficult to manage, leading to higher mortality rates, prolonged illness, long-term health complications, and rising healthcare costs.

The canonical model of resistance evolution is well established ^1^. Antibiotic exposure eliminates susceptible cells, whereas cells that acquire resistance mutations or genes survive, transmit these determinants stably to their progeny, and expand clonally. As a result of this selective sweep, these genetic determinants are uniformly shared across the population. Owing to this uniformity, the resistance of the entire population is summarized by a single metric—the minimum inhibitory concentration (MIC) ^2^. Based on the MIC-based testing, resistance is framed as a binary trait: an isolate either exhibits it or does not.

Yet growing evidence reveals a limitation of this paradigm. Many clinical isolates are neither fully resistant nor fully susceptible; instead they exhibit a subtle phenotype in which a minority of resistant cells remains embedded within a susceptible majority in a clonal population, a phenomenon clinically recognized as heteroresistance (HR) ^3,4^. A major challenge is that such isolates are not readily recognized by routine diagnostic tests yet still give rise to treatment failure, creating a discrepancy between diagnosis and therapeutic outcome ^3–6^.

HR was reported as early as the 1940s, during the early antibiotic era ^7,8^ and has since been observed across diverse bacterial species and antibiotic classes ^5^. Epidemiological surveys report that the incidence of HR has increased steadily with antibiotic use ^9^. In some settings, HR surpassed classical resistance as the predominant form of drug resistance ^10^. Analogous forms of HR and associated treatment failures have also been reported in fungal pathogens and cancer cells ^11,12^.

Despite its clinical prevalence and significance, the evolutionary origin of HR remains poorly understood. Antibiotic landscapes and strain backgrounds in clinical settings are complex, making it difficult to identify the selective conditions that favor HR. Although bet-hedging can evolve through modifications of preexisting stochastic mechanisms ^13,14^, its de novo evolution is generally considered challenging as various environmental constraints and fitness trade-offs severely restrict the relevant parameter regime ^15,16^. One proposed mechanism of HR involves transient gene duplication arising as a byproduct of DNA replication ^17^, which suggests that HR may be an intrinsic and preexisting property. On the other hand, a recent epidemiological study indicates that the incidence of HR has risen steadily with antibiotic use ^9^, suggesting that its emergence may be shaped by antibiotic selection. Currently, the evolutionary mechanism of HR remains unresolved.

To address this knowledge gap, we performed de novo evolution experiments starting from a fully susceptible *Escherichia coli* K-12 strain. By systematically varying antibiotic exposure regimes, we evolved HR and classical population-wide resistance from the same strain background, thereby identifying the selective conditions that drive these divergent evolutionary outcomes. We found that HR evolution was cryptic yet rapid and reproducible, underscoring a major diagnostic blind spot. Whole-genome sequencing and genetic reconstruction show low evolutionary barriers to HR. Direct head-to-head fitness comparisons of evolved strains revealed the adaptive basis of HR: it exploits phenotypic flexibility to enhance survival during antibiotic exposure while preserving growth during exposure gaps, unlike classical resistance, which is genetically locked into fixed fitness costs. We also discovered that clinical HR isolates exhibited genetic and fitness signatures similar to those of laboratory-evolved strains, indicating shared selective pressures. Finally, quantitative modeling unifies these findings into a predictive framework that accounts for distinct evolutionary progression of antibiotic resistance. Our findings provide fundamental insights into when and how HR emerges and why it is prevalent.

## Results

### Experimental reconstruction of divergent evolutionary routes to heteroresistance and classical resistance

Colistin is a critical last-line antibiotic for treating multidrug-resistant Gram-negative infections when few alternatives remain. Colistin-HR and associated antibiotic failure were first described in clinical isolates of *Enterobacter cloacae* ^4^ and have been reported in a wide range of Gram-negative pathogens (Supplementary Table 1). A recent survey indicates that HR surpassed classical resistance as the predominant form of colistin resistance ^10^. Based on this clinical importance and prevalence, we used colistin as a model antibiotic to investigate the evolutionary principles of HR. To assess whether our findings extend beyond colistin, we further repeated the evolution experiments using the β-lactam antibiotic meropenem.

Although HR isolates have been frequently found clinically, their genetic backgrounds are poorly defined, their evolutionary ancestors and intermediate states are unavailable, and their selective histories remain unknown. To experimentally reconstruct how HR emerges de novo, we performed controlled evolution experiments using *Escherichia coli* K-12, a genetically well-characterized and experimentally tractable model system.

In clinical laboratories, antibiotic susceptibility is most commonly assessed using inhibition-zone-based assays such as disk diffusion (Kirby–Bauer) and gradient diffusion (Etest) ^18^. Etest and disk diffusion performed on the ancestral *E. coli* strain produced a sharp inhibition boundary, yielding a single, well-defined MIC (Fig. 1a and Supplementary Fig. 1a-c). This MIC was well below the clinical breakpoint for colistin (4 µg/mL), indicating that the wild-type (WT) ancestral strain is fully susceptible to colistin as expected.

**Fig. 1.**
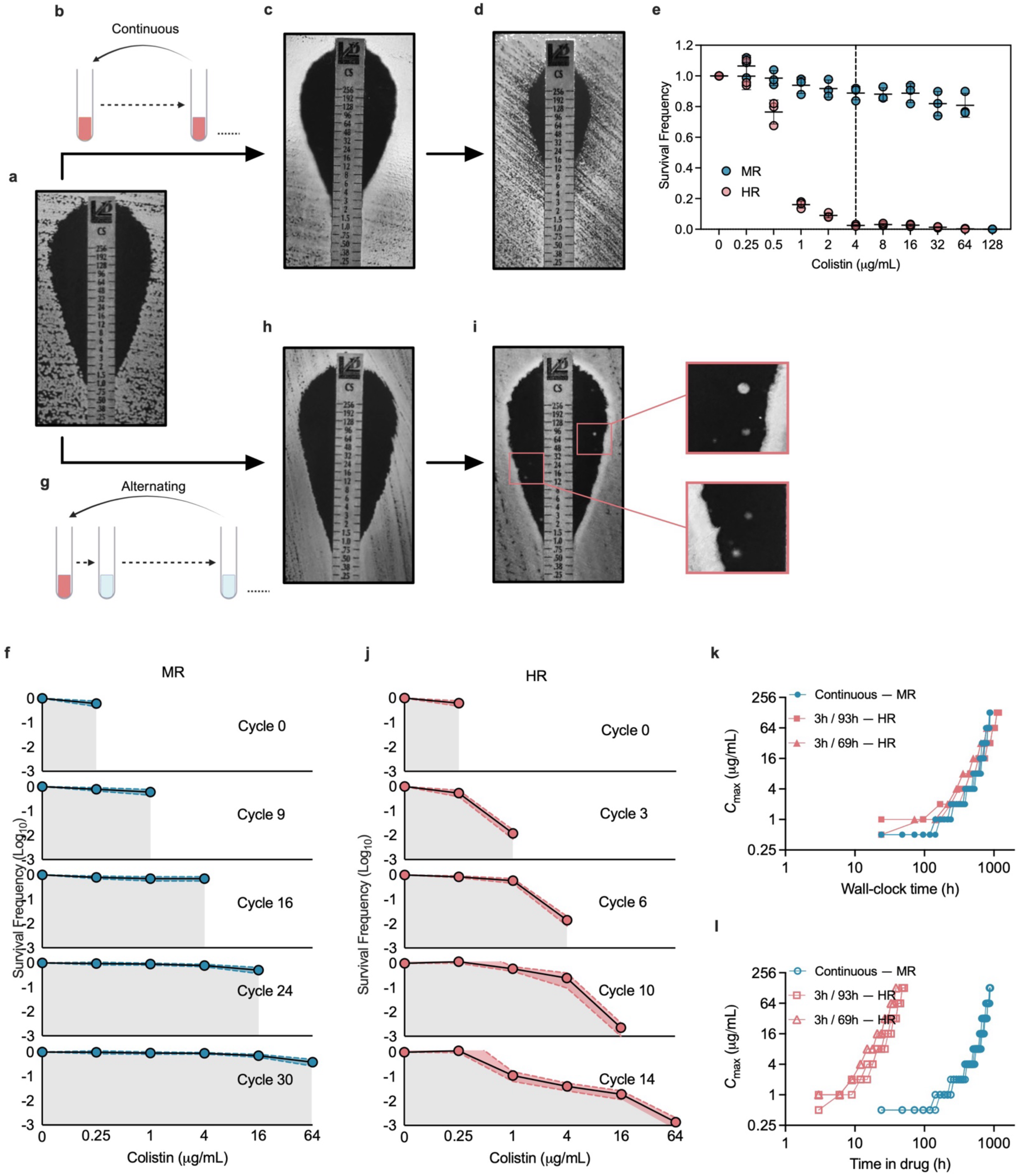
De novo evolution of heteroresistance (HR). (a) Representative inhibition-zone image of WT *E. coli* ancestral strain. See Supplementary Fig.1 for corresponding disk diffusion and PAP curve. (b) Schematic of the continuous-treatment regimen. One cycle consisted of 24 hours of treatment. In each cycle, cultures were exposed to both 0.5× and 1× MIC. When growth was observed at the latter, the antibiotic concentration was increased two-fold in the subsequent cycle. (c, d) Representative inhibition-zone images of mutants evolved under continuous selection. Mutants from Cycle 16 (c) and 30 (d) were shown. Clear inhibition boundaries progressively shift inward, consistent with increasing resistance. Disk diffusion assays show similar shifts (Supplementary Fig. 1a). (e) Population analysis profiling (PAP) curves of a WT (green color), the MR mutant (blue color) and HR mutant (red color). Final evolved strains (Cycle 30 for MR and 14 for HR) were analyzed. The MR strain maintains high survival across increasing drug concentrations, while the HR strain exhibits a gradual decline with a long high-concentration tail, reflecting survival of resistant minorities. The dashed line indicates the colistin clinical breakpoint (4 µg/mL). Three biological replicates (symbols) were tested; black lines and error bars denote mean ± s.d. (f) Evolutionary trajectories of MR resolved by PAP. Under continuous exposure, PAP curves shift uniformly rightward across cycles while retaining a sharp cutoff, indicating population-wide increases in resistance (MR). (g) Schematic of the alternating-treatment regimen. Each cycle consisted of 3 h of colistin exposure followed by 93 h drug-free recovery (total cycle length = 96 h). (h, i) Representative inhibition-zone images of mutants evolved under alternating exposure (T_off_ = 93 h). Mutants from Cycle 6 (h) and Cycle 14 (i) are shown. The primary inhibition boundary remains largely unchanged. At later stages, small colonies appear within the inhibition zone (insets). We made a similar observation for T_off_ = 69 h (Supplementary Fig. 3). (j) Under alternating exposure, PAP curves progressively develop a graded high-concentration tail across cycles, indicating expansion of increasingly resistant but progressively rarer subpopulations (HR). PAP curves are plotted on a semi-log scale to visualize low-frequency survivors. Solid dots represent the mean survival frequency of independent colonies (n=50) from five parallel cultures; shaded regions indicate the full range. We made a similar observation for T_off_ = 69 h (Supplementary Fig. 3). (k, l) Kinetics of resistance acquisition. Maximum resistant concentration, C_max_, as a function of wall-clock time (k) and cumulative time spent in drug (l), highlighting more rapid emergence of HR per unit drug exposure compared to MR. Symbols denote three replicate populations; lines connect individual data points where shown.

To accelerate evolution, the DNA mismatch repair gene *mutS* was deleted—this deletion increases mutation rates by ∼ 100-fold and is widely used in other evolutionary studies ^19,20^. Deletion of *mutS* had no detectable effect on antibiotic susceptibility (Supplementary Fig. 1b, c). All genetic reconstruction conducted later used the WT background with an intact *mutS* gene.

### Classical resistance evolves under continuous exposure or short-term fluctuations

We first followed a conventional evolution protocol in which cells were exposed continuously to antibiotic (Fig. 1b) ^21,22^. Selection pressure was ratcheted upward by increasing antibiotic concentration over time (Methods). Etest and disk diffusion assays showed a uniformly reduced inhibition zone that was displaced toward higher antibiotic concentrations relative to the WT ancestor (Fig. 1a, c, d; Supplementary Fig. 1a), indicating the evolution of resistance. The inhibition boundary remained abrupt, yielding elevated yet well-defined MICs.

To measure resistance quantitatively, we performed population analysis profiling (PAP). Although labor-intensive and therefore rarely used clinically, PAP quantifies cell survival as a function of antibiotic concentration, providing a direct readout of resistance frequency ^3,23,24^. A PAP curve of final evolved clones exhibited uniformly high survival across increasing concentrations, followed by an abrupt collapse at a high drug level (blue, Fig. 1e). This sharp cutoff lies well above the clinical breakpoint (dashed line) and therefore classifies the mutant as resistant. Importantly, this step-like profile indicates uniform survival up to a common concentration threshold (i.e., MIC) and explains the clear inhibition edges observed in Etest and disk diffusion assays (Fig. 1c, d and Supplementary Fig. 1a). These data are collectively consistent with the classical picture of homogeneous, population-wide antibiotic resistance. Hereafter, we refer to this phenotype as mono-resistance (MR).

To quantitatively resolve the underlying evolutionary dynamics, we performed PAP on isolates sampled across multiple stages of evolution. Across successive cycles, PAP curves consistently displayed a step-like profile, while the entire curve shifted progressively rightward (Fig. 1f). This shift indicates a rising MIC and a population-wide increase in resistance. Thus, continuous antibiotic exposure drives the progressive evolution of MR.

While continuous antibiotic exposure is commonly used to study resistance evolution, clinical antibiotic treatments are inherently dynamic ^25^. Scheduled dosing (e.g., oral or intravenous administration at fixed intervals) generates repeated fluctuations in antibiotic concentration over time. We simulated this fluctuating antibiotic landscape by exposing *E. coli* to alternating cycles of colistin treatment for a period of T_on_ and colistin-free recovery for T_off_. Previous studies have shown that cyclical regimens can increase antibiotic ‘tolerance’ ^26–29^, in which bacterial populations transiently halt growth to survive drug exposure. These studies were specifically designed to enrich tolerance by suddenly exposing stationary phase cells to very high drug concentrations (10 ∼ 500 × MIC), thereby killing all actively growing cells. Such a procedure was chosen because tolerance is defined by a dormant state ^30^. By contrast, resistance is defined by the ability of bacteria to continue growing in the presence of antibiotics ^30^. We therefore maintained cultures in exponential phase throughout evolution experiments by serial dilution.

We first implemented intermittent exposure with equal treatment and recovery periods (T_on_/T_off_ = 24/24 h, Supplementary Fig. 2a). Under these conditions, PAP curves again exhibited step-like profiles (Supplementary Fig. 2b). We then decreased T_on_ while extending T_off_ (12/36 h or 3/21 h, Supplementary Fig. 2a); yet PAP revealed the same step-like pattern (Supplementary Fig. 2a-d). Thus, continuous antibiotic exposure or short-term fluctuations result in evolution of MR.

### HR evolves under intermittent exposure with extended recovery periods

In clinical treatment settings, extended antibiotic-free intervals can arise for multiple reasons, including non-adherence to prescribed regimens (e.g., missed doses) ^31^ and repeated treatment episodes following infection relapse ^32^. In addition, antibiotic penetration can vary substantially across tissue compartments ^33^; as bacteria move between compartments, they may experience effective gaps in antibiotic exposure. Transmission of bacteria between hosts who are being treated and untreated may create even longer periods of intermittent exposure ^34,35^.

Motivated by these considerations, we next extended the antibiotic-free interval in our evolution experiments to multiple days. We kept T_on_ at 3h, the minimum duration required to reliably eliminate susceptible cells (Supplementary Fig. 1d) and extended T_off_ to 69 h and 93 h, generating 3-day and 4-day treatment cycles, respectively (Fig. 1g and Supplementary Fig. 3a). Although the length of each cycle (T_on_ + T_off_) was substantially longer than in the regimens described above, the total duration of evolution was kept comparable (∼ 1 month).

Under this regime, we observed a distinct evolutionary outcome. Etest and disk diffusion assays produced inhibition zones nearly indistinguishable from those of the susceptible ancestor, indicating that the apparent MIC did not change (Fig. 1h, i; Supplementary Fig. 1a and 3). To quantify cell survival, we conducted PAP on the final evolved isolates. In contrast to the step-like profile seen for MR, the PAP curve declined gradually with increasing drug concentration (Fig. 1e, red). Survival fell to very low levels at concentrations below the clinical breakpoint (dashed line). Therefore, the majority of cells remained susceptible despite the long duration of the evolution experiment.

Yet the PAP curve also displayed a pronounced long tail, with low but nonzero survival across a broad range of concentrations far above the apparent MIC (Fig. 1e, red). Motivated by this observation, we closely re-examined the Etest and disk diffusion plates and detected distinct colonies within the primary inhibition zone in late-cycle isolates (Fig. 1i, inset; Supplementary Fig. 1a and 3). Together, these data reveal rare but highly resistant cells embedded within an otherwise susceptible population, i.e., heteroresistance (HR). Thus, intermittent exposure with extended antibiotic-free recovery periods drives HR evolution.

To quantitatively resolve how a susceptible WT strain developed HR under this condition, we performed PAP on isolates sampled across multiple stages, plotting survival frequency on a semi-log scale to resolve low-frequency survivors (Fig. 1j and Supplementary Fig. 3). Over successive cycles, survivors appeared at progressively higher colistin concentrations but at low frequencies, producing graded tails in the PAP curve. These tails widened over time, expanding the concentration range over which rare cells survived. This trajectory contrasts sharply with population-wide resistance observed in the MR evolution above (Fig. 1f).

Broadly, HR refers to the presence, within an otherwise susceptible isolate, of a minor subpopulation capable of surviving antibiotic concentrations at or above the clinical breakpoint ^24^. More recently, a quantitative criterion has been proposed to classify an isolate as HR if it contains subpopulations (frequency ≥ 10^−6^ – 10^−7^) capable of growth at ≥ 4 or 8× the MIC of the main susceptible population ^3,23,24^. Our evolved isolates meet this quantitative criterion for HR already at early evolutionary stages (e.g., Cycle 6 in Fig. 1j).

We next repeated the experiment using meropenem. The results mirrored those observed with colistin: continuous meropenem exposure or short fluctuations selected for MR (Supplementary Fig. 4a-c), while intermittent exposure with long antibiotic-free recovery periods favored HR (Supplementary Fig. 4d, e). This result again indicates that the temporal structure of antibiotic exposure governs the evolutionary divergence between MR and HR.

### HR emerges cryptically, reproducibly, and rapidly

Notably, at the early stages (e.g., Cycle 6), the HR phenotype remains undetectable by Etest or disk diffusion, which yield inhibition boundaries identical to the susceptible ancestor and lacked obvious colonies within the zone (Fig. 1h; Supplementary Fig. 1a and 3b, c). Such colonies appeared only at later evolutionary cycles (Fig. 1i; Supplementary Fig. 1a and 3b, c). Thus, HR emerges as a cryptic phenotype that escapes routine MIC-based detection.

Here, the PAP curves (Fig. 1f, 1j) were obtained by analyzing 50 independent colonies isolated from five parallel evolution cultures (10 colonies per culture). Each PAP curve was obtained from a single, purified clonal isolate; thus, the gradual decline in survival frequency (Fig. 1j) reflects intrinsic phenotypic heterogeneity within a clonal population rather than a mixture of distinct genotypes. Importantly, all 50 PAP curves closely overlapped (shaded regions, with symbols indicating the mean). These results show that HR evolves robustly and reproducibly as a clonal phenotype.

We next analyzed the timescale of evolution. We quantified evolutionary progress using C_max_, defined as the highest colistin concentration up to which survival was detectable in PAP. When C_max_ is plotted against elapsed time, it increases on comparable wall-clock time in the continuous and intermittent regimens (Fig. 1k). However, the intermittent protocols expose cells to colistin for only a small fraction of each cycle (drug duty cycles of 1/24 or 1/32), whereas continuous exposure maintains drug pressure throughout. Accordingly, when C_max_ is plotted against cumulative antibiotic exposure time, C_max_ increased >10-fold faster in HR than MR (Fig. 1l). Thus, per unit antibiotic exposure, HR evolves far more efficiently than MR.

### Single mutations are sufficient to generate HR

The genetic mechanisms of classical resistance have been well studied ^36^. By contrast, HR exhibits distinct phenotypic characteristics, raising the question of whether it requires complex genetic changes. Our experiments evolved mutants from a well-defined genetic background (*E. coli* K-12) and followed their lineages, providing a unique opportunity to identify genetic signatures that enable HR. We performed whole-genome sequencing of HR-evolved strains, identifying approximately 15 mutations that have accumulated over the course of evolution (Supplementary Fig. 5a). To assess their contributions, each mutation was introduced separately into the WT background. Most mutations had no detectable or only limited effects on colistin resistance (Supplementary Fig. 5b). By contrast, two mutations—one in *pmrB* and one in *lpxC*—produced HR phenotypes (Fig. 2).

**Fig. 2.**
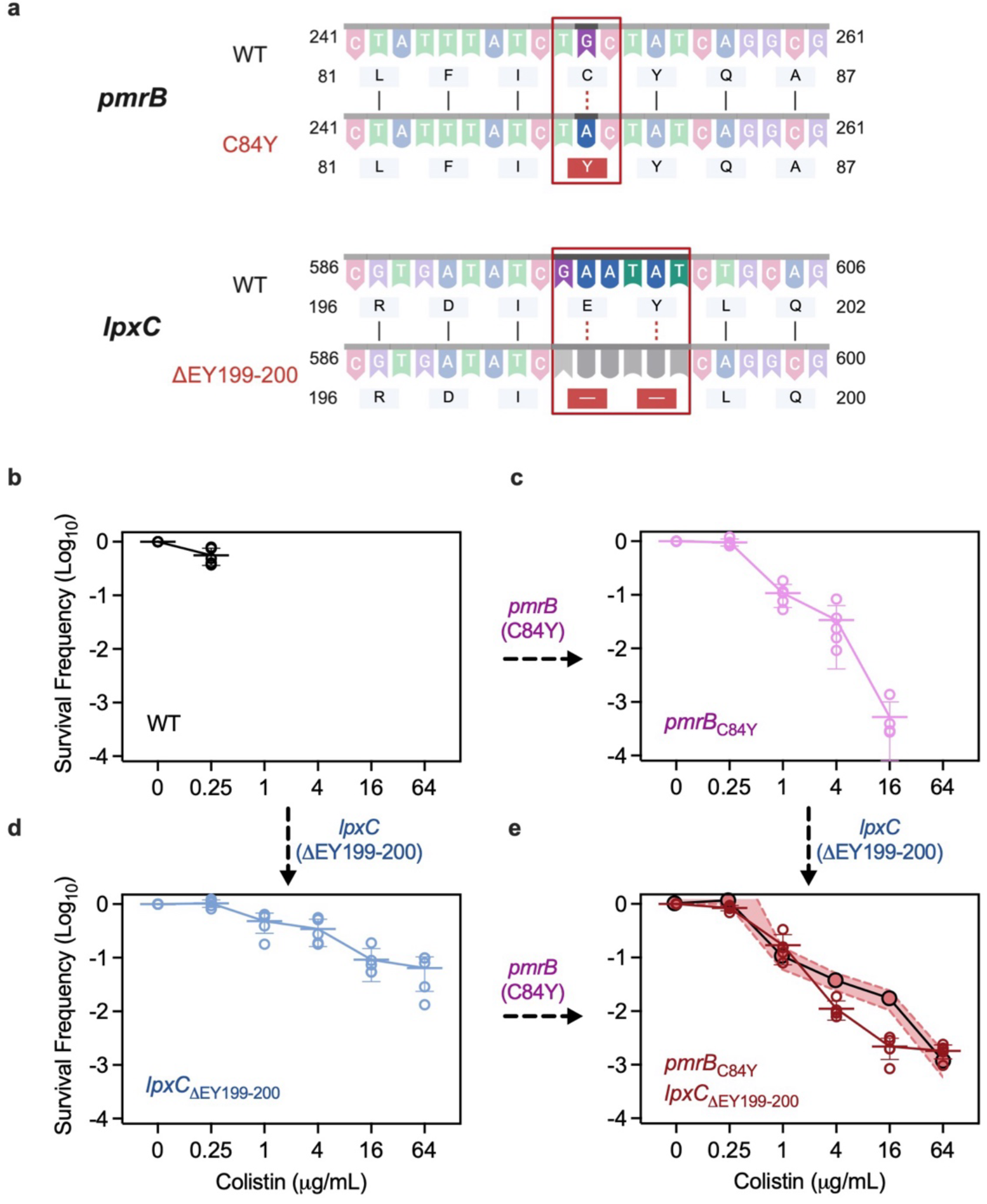
Genetic analysis of evolved HR strains. (a) Colistin-evolved HR strains were analyzed by whole-genome sequencing (Supplementary Fig. 5a). Each mutation identified was introduced individually into a WT background using CRISPR-assisted λ-Red recombination (Supplementary Fig. 5b), identifying two critical mutations — the *pmrB* (C84Y) and *lpxC* (ΔEY199–200). (b-d) Introduction of either *pmrB*(C84Y) or *lpxC*(ΔEY199–200) into WT was sufficient to produce HR phenotypes. Survival frequency is shown on a log scale. Different symbols represent independent replicate cultures (n = 5); Lines and error bars indicate their mean and standard deviation. (e) Combining these two mutations (red line with symbols) recapitulated the PAP profile of the evolved HR strain (Cycle 14, shaded black line).

The nucleotide changes underlying these mutations are shown in Fig. 2a. Colistin targets lipid A within lipopolysaccharide ^37,38^. Consistent with this mode of action, both mutations mapped to genes involved in lipid A pathways. *pmrB* encodes the sensor kinase component of the PmrAB two-component regulatory system, which regulates lipid A modification ^39^. We identified a missense mutation (C84Y) within the N-terminal transmembrane/sensory region previously characterized for signal detection (Fig. 2a) ^40^.

*lpxC* encodes an essential enzyme catalyzing the first committed step in lipid A biosynthesis ^41^. Because *lpxC* is essential, it cannot be deleted without loss of viability; the fact that the evolved strain remains viable indicates that the mutant protein retains function. We found a two-residue deletion (ΔEY199–200, Fig. 2a) within a hydrophobic pocket implicated in substrate positioning, adjacent to the essential catalytic core ^42^.

Because transient gene amplification was proposed as a potential mechanism for HR ^17^, we additionally examined copy-number variation in the evolved strains. Read-depth analysis occasionally yielded modestly elevated copy-number estimates (∼2–3×) at a few isolated loci (Extended Data). This magnitude is far smaller than that of previously reported amplifications (often tens-to hundreds-fold) ^17,43^. Moreover, genes immediately adjacent to these loci retained baseline coverage (i.e., single-copy), lacking the contiguous multi-kilobase amplification structure expected for bona fide segmental duplication ^17,44^ (Extended Data). Together, these analyses argue against segmental amplification as the primary basis of HR in our evolved strains.

Importantly, introduction of either *pmrB* or *lpxC* mutation alone into the susceptible WT background was sufficient to convert the PAP profile to HR (Fig. 2b-d). The combined *pmrB* and *lpxC* mutations recapitulated the PAP shape of the evolved HR strain (Fig. 2e). These results show that HR does not require complex genomic remodeling, but can arise from single mutations. This low genetic barrier helps explain why HR evolves rapidly and reproducibly as observed above.

### HR maintains its adaptive advantage through phenotypic partitioning and dynamic enrichment

The rapid and reproducible emergence of HR in our evolution experiments suggests that this phenotype confers a selective advantage. To identify this advantage, we leveraged our experimental design, which yielded both HR and MR from the same parental background. We compared the HR mutant from Cycle 10 and the MR mutant from Cycle 24, because both were able to grow in colistin up to the same maximal concentration (*C*_max_=16 µg/mL). This allows a direct fitness comparison between HR and MR at matched resistance levels.

Given that the parent strain fails to grow at colistin concentrations above 0.25 µg/mL (Supplementary Fig. 1), the ability of MR and HR strains to survive up to 16 µg/mL, which is well above the clinical breakpoint 4 µg/mL, highlights the benefit of resistance. Yet this is accompanied by a cost; under antibiotic-free conditions, both MR and HR strains exhibited reduced growth rates relative to their susceptible ancestor (Supplementary Fig. 6a). By introducing the *pmrB*(C84Y) and *lpxC*(ΔEY199–200) mutations into WT (Fig. 2a), we found that these two mutations recapitulated the fitness cost of the HR strain (Supplementary Fig. 7).

However, when we directly compared the HR and MR strains, the former grew faster than the latter (Fig. 3a, left panel); see Supplementary Fig. 6a for their growth curves, and Supplementary Fig. 4f for meropenem-HR and MR. We additionally confirmed faster growth of HR than MR consistently throughout the evolution (Supplementary Fig. 6b and 4g). Thus, although resistance incurs a fitness cost, this cost is substantially lower in HR than in MR.

**Fig. 3.**
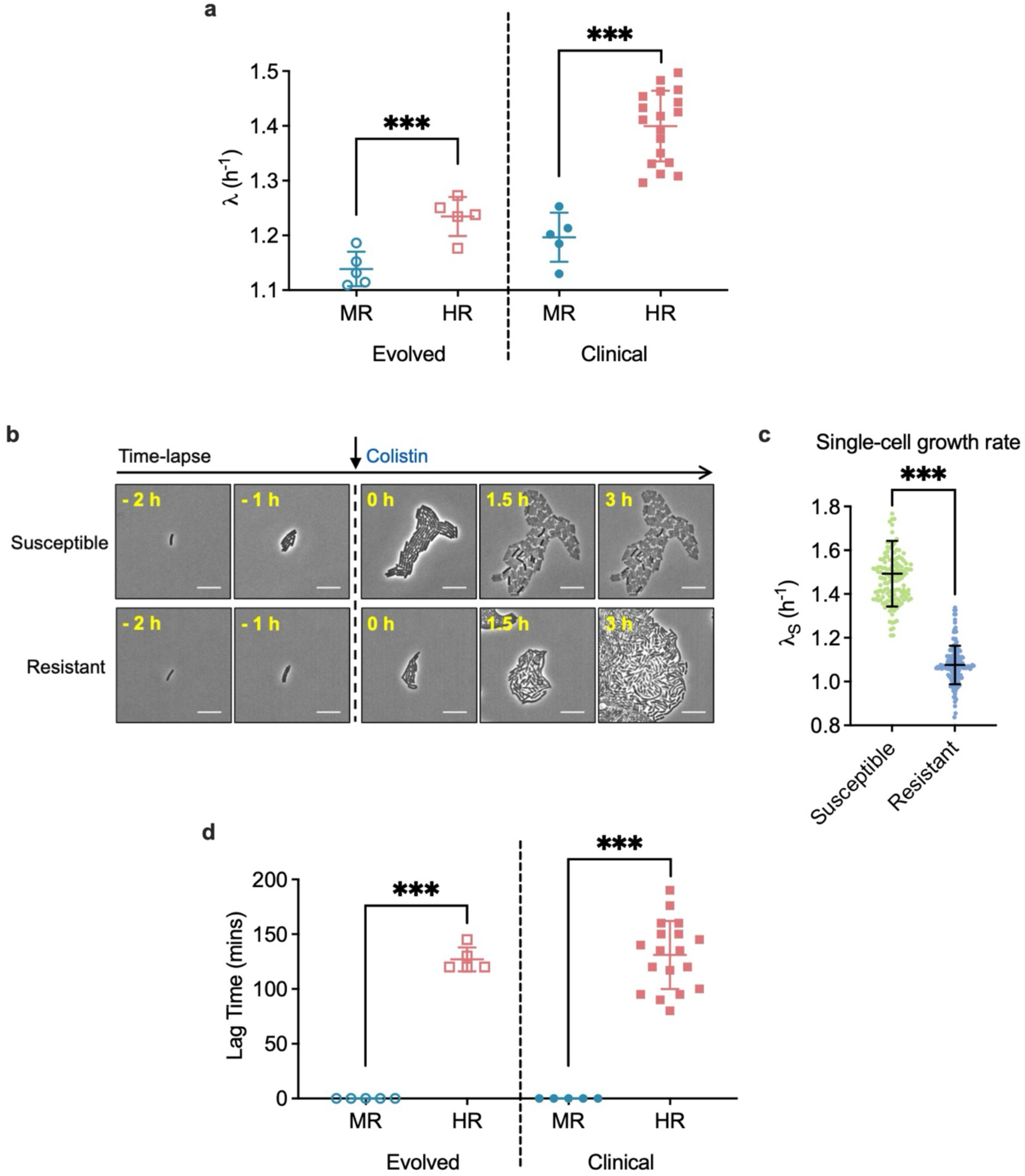
Fitness analysis of HR and MR. (a) Left panel: Growth rates (λ) of evolved strains. The MR strain from Cycle 24 and the HR strain from Cycle 10 was characterized. Different symbols represent independent replicate cultures (n = 5). Right panel: Growth rate of clinical isolates collected through the CDC’s Multi-site Gram-negative Surveillance Initiative (MuGSI) ^10^. The dataset includes colistin-MR (n = 5) and HR (n = 18) isolates. All isolates belong to the *Enterobacterales* order and comprise predominantly *Enterobacter*, *Escherichia*, and *Klebsiella* species. Species identities and individual growth rates are provided in Source Data. (b) Time-lapse microscopy of an evolved colistin-HR strain. Images to the left of the arrow show the pre-treatment phase; those to the right show the post-treatment phase. Cells were classified based on their response to colistin post-treatment: resistant (surviving at 16 µg/mL) or susceptible (killed at 2 µg/mL). Scale bar: 10 µm. (c) At single-cell level, susceptible cells exhibited significantly higher growth rates prior to colistin exposure. Three independent experiments were conducted, with 50 cells per group per replicate. (d) Lag times of lab-evolved strains (left panel, n = 5) and clinical isolates (right panel, sample sizes were n = 5 for MR and 18 for HR) after exposure to antibiotic at 0.5× the strain-specific MIC. Black lines and error bars indicate the mean and standard deviation. ***p < 0.001, based on one-way ANOVA with multiple comparisons.

**Fig. 4.**
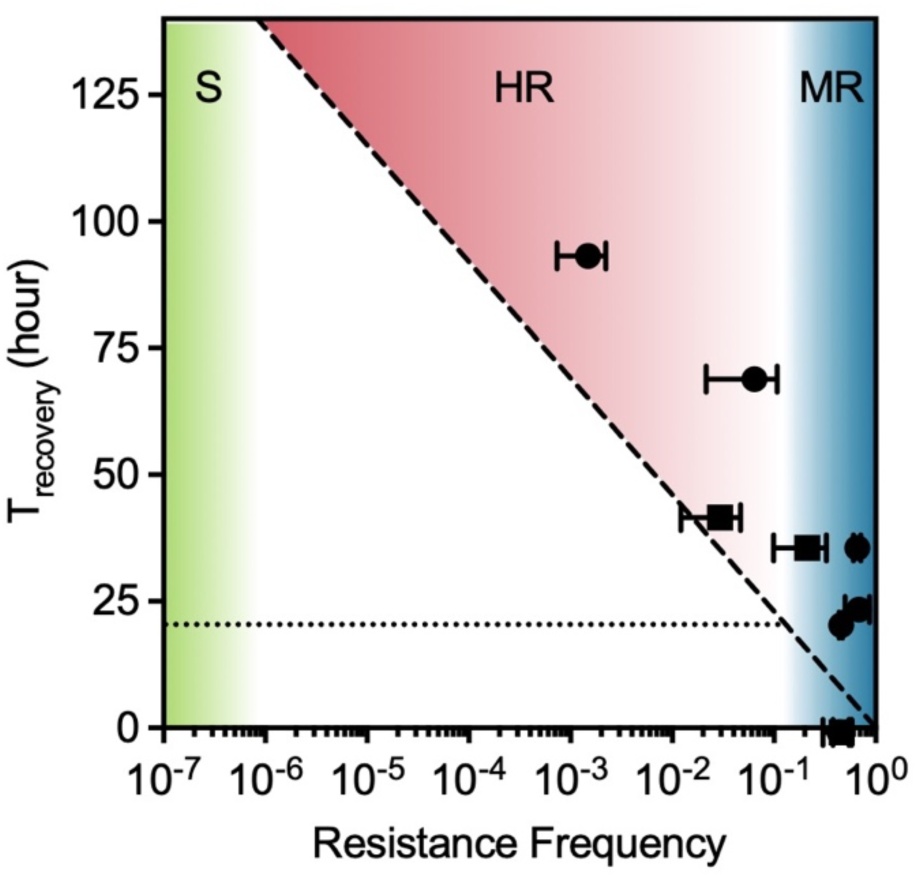
Phase diagram predicting when HR is evolutionarily viable. The dashed line (Eq. 7) indicates the minimum T_recovery_ required for a population with a given resistant-subpopulation frequency to emerge. Shaded regions denote operational classifications of susceptibility (S), HR, and MR. The red region indicates the HR regime. Black circles (colistin) and squares (meropenem) show experimental evolution outcomes across distinct schedules; error bars indicate standard deviation. Experimental points lie close to, but above, the predicted boundary, supporting the model.

To understand the basis of HR’s reduced fitness cost, we analyzed growth at single-cell resolution. The antibiotic responses of individual cells reveal heterogeneous phenotypes within the HR population. Cells were first grown exponentially under antibiotic-free conditions (left of the arrow in Fig. 3b) and then exposed to colistin (arrow). Many cells were killed upon exposure to a colistin concentration (2 µg/mL; top panel, right of the arrow in Fig. 3b), indicating that they were susceptible. In contrast, a subset of cells continued to grow even after exposure to a high colistin concentration (16 µg/mL; bottom panel) and were classified as resistant.

After this classification, we quantified the antibiotic-free single-cell growth rate, *λ*_S_, by analyzing the time-lapse trajectories prior to antibiotic exposure. This analysis revealed a clear fitness trade-off. In the absence of antibiotics, resistant cells grew slowly (Fig. 3b, c), with growth rates comparable to those of MR (Fig. 3a, left panel). In contrast, susceptible cells grew fast, forming larger microcolonies prior to exposure (Fig. 3b) and exhibiting higher growth rates (Fig. 3c). We observed the same cell-to-cell heterogeneity in the meropenem-evolved HR strain (Supplementary Fig. 8). Together, these results show that a HR population comprises phenotypic states with distinct trade-offs, partitioning growth and survival across single cells.

We next analyzed how these phenotypes were managed within a population. Under antibiotic-free conditions, the fraction of resistant cells remains low (Supplementary Fig. 9a, b). Upon antibiotic exposure, this resistant subpopulation expands to comprise nearly the entire population, and then contracts back to its basal level once the antibiotic is removed (Supplementary Fig. 9a, b). This behavior contrasts sharply with MR, in which the resistant fraction remains fixed near 100% regardless of environmental conditions (Supplementary Fig. 9c, d).

Together, these observations indicate a dynamic strategy of HR: resistant phenotypes are transiently enriched during antibiotic exposure to ensure survival, while susceptible, fast-growing cells dominate in antibiotic-free environments to maximize population growth. This population-level restructuring of single-cell states provides a mechanistic basis for how HR achieves robust survival while maintaining a lower fitness cost of resistance.

### Clinical isolates exhibit genetic and fitness signatures similar to laboratory-evolved strains

The analyses above establish a genetic basis and fitness advantage of HR in lab-evolved strains. To evaluate the clinical relevance of these findings, we next examined clinical isolates. Unlike our lab strains, which were derived from a single genetic background under defined exposure regimens, clinical isolates span diverse genetic backgrounds and unknown treatment histories. We next asked whether the defining features of lab-evolved HR are conserved in clinical isolates.

First, we surveyed prior clinical studies of colistin HR in the literature, primarily from Gram-negative ESKAPEE pathogens, including *Enterobacter* spp., *Escherichia coli*, *Klebsiella pneumoniae*, *Acinetobacter baumannii*, and *Pseudomonas aeruginosa* (see Supplementary Table 2). When we compiled the reported mutations, we found that genes in the *pmr* and *lpx* pathways were among the most frequently mutated loci (∼50% and 15% respectively, Supplementary Table 2 and Supplementary Fig. 10).

We next measured the fitness of clinical HR and MR isolates. We utilized the U.S. CDC Emerging Infections Program ^10^, analyzing a collection comprising primarily *Enterobacter*, *Escherichia*, and *Klebsiella* species. The population-level growth measurements showed that HR isolates consistently grew faster than MR isolates (Fig. 3a, right panel). Therefore, a lower fitness cost of resistance is a robust feature of HR in clinical isolates as well.

The microscopy analyses further revealed a similar phenotypic trade-off between growth and resistance at the single-cell level in clinical isolates (Supplementary Fig. 11). Moreover, the size of the resistant subpopulation in HR strains dynamically shifted in response to environmental conditions (Supplementary Fig. 9e, f), in contrast to a clinical MR isolate in which the resistant fraction remains fixed near 100% regardless of environment (Supplementary Fig. 9g, h).

Together, our results from clinical isolates and lab-evolved strains converge across genetic, fitness, and single-cell phenotypes. This convergence, despite diverse genetic backgrounds, suggests that comparable selective pressures operate in clinical settings, similar to those imposed in our evolution experiments. Importantly, our observation of HR’s fitness advantage provides a mechanistic explanation for its widespread occurrence ^3,5^.

### Fitness modeling recapitulates the distinct selective regimes underlying HR and MR

These findings raise an important question: why is HR not always favored over MR? Although HR reduces the constitutive fitness cost of resistance during antibiotic-free periods, it carries a distinct vulnerability: only a small minority of cells are resistant at the onset of antibiotic exposure. Consistent with this limitation, HR populations experience a substantial lag phase upon antibiotic exposure, until minority resistant cells expand sufficiently to support population growth (Fig. 3d). Such prolonged lag phases were observed in both laboratory-evolved strains (left panel) and clinical isolates (right panel), revealing a severe deficit at treatment onset.

Integrating this advantage and limitation into a mathematical framework, we sought to quantitatively model and predict the evolutionary conditions that favor HR. In our evolved HR populations, the frequency of cells surviving the highest colistin concentrations tested, *C*_max_—that is, the resistant-subpopulation frequency (RF)—is typically ∼0.01–0.1% of the total population (Fig. 1j), corresponding to an approximately 10^3^–10^4^-fold initial deficit upon antibiotic exposure. In antibiotic-free conditions, however, HR populations grow faster than MR populations by ∼0.1 h ¹ (Fig. 3a). Thus, HR can offset an initial disadvantage at treatment onset through a growth advantage during recovery.

We quantified this cost-benefit balance by calculating the geometric mean fitness over one cycle consisting of an antibiotic exposure period (T_on_) followed by an antibiotic-free recovery period (T_recovery_) (Supplementary Text). Eq. 7 gives the minimum recovery time required for HR to compensate for its initial deficit: 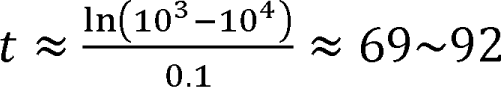 hours. Therefore, our model explains why long recovery periods were required to evolve HR, consistent with our experimental finding (T_recovery_ = 69 and 93 h).

However, our previous study has shown that the RF in clinical isolates can vary over several orders of magnitude ^45^. Using Eq. 7, we therefore calculated the T_recovery_ as a function of RF, yielding the diagonal dashed line in Fig. 4. This predicted line reveals that the rarer the resistant subpopulation, the longer the recovery time required for its growth advantage in drug-free conditions to compensate for its deficit during treatment. Importantly, this predicted line defines the minimum T_recovery_ needed for a given RF; HR is a competitive evolutionary outcome only above this boundary.

To test this theoretical result, we mapped our colistin and meropenem evolution outcomes onto this phase diagram. We calculated the RFs by the number of survivors at *C*_max_ in our evolved strains. Plotting the T_recovery_ used in each experiment against this frequency places experimental points consistently near, but above, the predicted minimum boundary (Fig. 4), supporting our model.

For practical interpretation, we overlaid standard operational classifications. At low frequencies (≲10 –10), resistant cells fall near the detection limit of PAP. In this regime, isolates are operationally treated as susceptible (S; green region in Fig. 4) ^3,23,24^. At the opposite extreme, when a macroscopic fraction of the population is resistant (e.g., ≳0.1–0.5), it is readily detected by routine assays and categorized as classical resistance (MR; blue region). The intermediate regime corresponds to HR (red region). Overlaying this classification with our model defines a phase diagram that unifies S, HR, and MR as distinct evolutionary regimes.

Importantly, this phase diagram shows that HR is favored only when recovery intervals are well above ∼20 h (dotted line), with longer recovery intervals supporting lower resistant-subpopulation frequencies. These intervals are substantially longer than the within-day fluctuations generated by standard dosing schedules. By contrast, such extended antibiotic-free gaps can readily arise from non-adherence to prescribed antibiotic regimens, including missed doses, which are common in outpatient settings ^31^. These results identify such treatment gaps as a key selective condition for HR evolution.

Notably, the relevant timescale depends on fitness parameters. In our framework (Eq. 7), the minimum recovery time required for HR scales inversely with the antibiotic-free growth advantage of HR over MR (Δλ). In laboratory-evolved strains, we measured Δλ ≍ 0.1 h^-1^ (Fig. 3a). In clinical isolates, the growth-rate advantage was approximately two-fold larger (Fig. 3a), which shifts the HR-favored boundary downward by roughly a factor of two. As a result, the recovery intervals required to favor HR can encroach upon within-day dosing timescales. Therefore, even standard dosing fluctuations may, under some conditions, contribute to HR evolution.

## Discussion

Evolution critically depends on history, yet the details of that history are often unknown, obscuring the adaptive origins of evolved traits ^46^. In clinical settings, antibiotic landscapes and strain backgrounds are inherently complex and poorly defined, making it difficult to delineate evolutionary determinants. As a result, it has remained unclear when and how HR emerges and why it is so prevalent. Our study addresses this gap by directly demonstrating the de novo evolution of HR.

We show that HR evolves rapidly and reproducibly under treatment interruptions. Such interruptions are not unique to antibacterial therapy but also often arise in antifungal and anticancer regimens ^31,47^. Thus, the selective mechanism uncovered here may extend well beyond antibacterial therapy.

This evolutionary pathway poses a particular clinical challenge. While resistance evolution may be unavoidable under sustained antibiotic selection, classical resistance (MR) is readily detectable by standard susceptibility testing, enabling rational therapeutic adjustment and stewardship responses. In contrast, we demonstrated that HR evolution is cryptic, obscuring both its emergence and its progression. Although we used PAP to detect HR in the present study, its labor-intensive and low-throughput nature limits routine clinical use ^3^. Consequently, HR creates a diagnostic blind spot that constrains real-time epidemiological surveillance and permits treatment failure to occur without a clear warning signal.

Why does such diagnostically elusive resistance emerge? Our fitness analyses provide a mechanistic explanation. By generating phenotypic heterogeneity, a HR population minimizes the fitness costs associated with resistance during drug-free intervals. Upon antibiotic exposure, rare resistant cells enable population survival. Their frequency dynamically shifts, decreasing to low levels in antibiotic-free conditions. Thus, the cryptic nature of HR is not directly selected per se; instead, it arises as a byproduct of the adaptive population structure. Once established, the low fitness burden of HR facilitates its maintenance and spread.

In summary, our study reveals how the temporal structure of antibiotic exposure determines the evolutionary pathway of resistance, defining the selective regime that favors HR. Our results establish HR as a readily evolvable adaptive strategy under treatment interruptions, in which phenotypic heterogeneity enables antibiotic survival with minimal fitness cost, thereby explaining its emergence and widespread occurrence.

## Methods

### Bacterial strains and culture conditions

*Escherichia coli* K-12 strain NCM3722 (referred to as NMK1) was used. A Δ*mutS* derivative of NMK1 (designated NMK476) was constructed via P1 transduction from the Keio collection. *Enterobacter cloacae* clinical isolates were collected through the Georgia Emerging Infections Program, as part of the CDC’s Multi-site Gram-negative Surveillance Initiative (MuGSI) in Georgia, USA.

Luria Broth (LB; Fisher Scientific #BP1426500) and LB agar (Fisher Scientific #BP1425500) were used for cell culturing. 5mL of cultures were incubated in test tubes with shaking at 250 rpm at 37 °C. OD_600_ readings were taken every 20–35 minutes using a Genesys20 spectrophotometer.

### Antibiotics

Colistin sulfate (#1264-72-8) was obtained from Sigma-Aldrich. Meropenem trihydrate (#119478-56-7) was obtained from Research Product Industry. Ampicillin (#69-52-3), kanamycin sulfate (#70560-51-9) and chloramphenicol (#56-75-7) were purchased from Bio Basic.

### Experimental evolution assays

Experimental evolution assays were conducted using NCM3722 Δ*mutS* (NMK476). A single colony was picked from an LB agar plate (previously streaked from -80°C glycerol stocks) and cultured overnight for 16 hours in LB broth with shaking at 250 rpm.

#### Continuous exposure

Overnight cultures were diluted into fresh LB medium containing either 0.5× MIC (e.g., 0.25 µg/mL) or 1× MIC (e.g., 0.5 µg/mL) of an antibiotic to a starting OD_600_ of 0.01. Cultures were incubated at 37 °C with constant shaking for 24 hours (one cycle). Bacterial growth was assessed visually and/or by measuring OD_600_. If growth was observed only in the 0.5× MIC condition, cultures were again diluted into fresh 0.5× and 1× MIC media, and the cycle was repeated. Once growth was detected in the 1× MIC condition, this culture was diluted into LB medium containing 1× and 2× MIC concentrations (e.g., 0.5 and 1 µg/mL). This stepwise two-fold increase in drug concentration continued every 24 hours, with cultures passaged to higher concentrations only after successful growth at the latter concentration. Throughout the experiments, cultures were diluted into fresh LB with drug at OD_600_ = 0.01 every 12 hours.

#### Alternation of antibiotic and antibiotic-free phases

We first conducted treatment cycles of 24 hours of antibiotic exposure and 24 hours of antibiotic-free phase (24 hr on/24 h/off). Cultures were initially diluted from overnight growth into fresh LB containing 0.5× MIC (e.g., 0.25 µg/mL) and 1× MIC (0.5 µg/mL) of an antibiotic at a starting OD_600_ of 0.01 and incubated at 37 °C with shaking for 24 hours. If growth occurred only in the 0.5× MIC condition, cultures from this condition were washed twice with fresh LB to remove residual antibiotic, then transferred into drug-free LB at an OD_600_ of 0.001 and incubated for 24 hours. Cultures were then diluted back into fresh LB containing 0.5× and 1× MIC drug concentrations at OD_600_ = 0.01, and the cycle was repeated.

Once growth was observed in the 1× MIC tube, cells from that culture were again washed, transferred to drug-free LB at OD_600_ = 0.001, incubated for 24 hours, and exposed to 1× and 2× MIC. These alternating cycles were repeated with stepwise increases in drug concentration.

We tested different cycling periods as illustrated in the figures (Fig. 1 and Supplementary Figs. 2-4). During drug-free phases, cultures were diluted into fresh LB at OD_600_ = 0.001 every 12 hours.

### Population analysis profile (PAP) tests

The PAP assay enumerates colony-forming units (CFU) on plates containing increasing drug levels. During evolution experiments, cultures were streaked on agar plates. Single colonies were resuspended in LB broth to OD_600_ ∼ 0.1, and serially diluted in LB in a 96-well plate (Corning #3598). 5 µL from each dilution was plated on LB agar containing antibiotics at indicated concentrations. CFU on drug-containing plates normalized by CFU on drug-free plates (after 24 hours of incubation) was calculated to determine resistance frequency.

### Etest and disk diffusion

Bacterial cultures were harvested during exponential phase, centrifuged at 8000 × g for 3 min, and resuspended in 1× PBS. Cell suspensions were adjusted to 0.5 McFarland standard. Sterile cotton swabs were dipped into the cell suspension and then used to evenly swab the surface of Mueller–Hinton agar plates. For Etest, colistin MIC test strips (0.016–256 μg/mL) (Liofilchem™ #921411) were placed according to the manufacturer’s instructions. For disk diffusion, sterile 6-mm disks were loaded with 64 μg of colistin sulfate and applied to the agar surface. Plates were incubated at 37 °C for 18–20 h and imaged after incubation.

### Whole-genome sequencing

Genomic DNA was extracted using the QIAamp DNA Mini Kit (Qiagen #51304) according to the manufacturer’s instructions. Purified DNA samples were submitted to SeqCenter (Pittsburgh, PA, USA) for whole-genome sequencing and bioinformatic analysis.

### Strain construction: genome editing

A CRISPR-assisted λ-Red recombination system ^48^ was used to introduce precise mutations into the genome of wild-type *E.coli* NCM3722 (NMK1). The λ-Red recombinase (expressed by pKD46) and CRISPR-Cas9 system were co-utilized to promote homology-directed repair at the targeted genomic site. A 20-bp spacer (listed in Supplementary Table 3) was cloned into the CRISPR plasmid (pCRISPR) using BsaI digestion and T4 DNA ligase.

Electrocompetent NMK1 cells carrying pKD46 (NMK99) were prepared by growing cultures at 30 °C to an OD of 0.2–0.3, inducing λ-Red expression with 0.2% L-arabinose, and washing cells sequentially with ice-cold water and 10% glycerol.

For co-electroporation, 50 µL of electrocompetent cells were mixed with 100 ng each of pCas9 and pCRISPR::spacer plasmids, along with 200 pmol of single-stranded DNA repair template (60 nt) (listed in Supplementary Table 3). Electroporation was performed at 1.8 kV, 25 µF, and 200 Ω, followed by recovery in SOC medium at 37 °C for 2 h. Transformants were selected on LB agar containing ampicillin (100 µg/mL), kanamycin (50 µg/mL), and chloramphenicol (25 µg/mL), and incubated at 30 °C overnight.

Genomic DNA from selected colonies was isolated, and the targeted region (∼300 bp flanking the mutation site) was amplified by PCR and verified via Sanger sequencing to confirm the desired mutation. For plasmid curing, verified mutants were propagated without antibiotics and incubated at 42 °C to eliminate the temperature-sensitive pKD46.

### Microscopy

Cultures grown to OD_600_ ∼0.1 were placed on a 35 mm glass-bottom dish (Cellvis) and covered with a 1.5% LB-agarose gel pad. Imaging was performed using an Olympus IX83 inverted microscope equipped with a 60× oil immersion phase-contrast objective within a pre-warmed (37 °C) incubation chamber (InVivo Scientific).

Time-lapse imaging of cells for growth rate determination was performed over a 2-hour period with an interval time of 10 minutes before exposure to colistin or meropenem. To assess the viability of growth-rate tracked cells, imaging was continued every 30 minutes following application of 2 µg/mL or 16 µg/mL colistin or meropenem to the gel pad.

Images were segmented and cell area was measured using the MicrobeJ plugin for Fiji/ImageJ (version 5.13 I). Growth rates were calculated from areal changes in 150 cells per subpopulation.

## Statistical analyses

Statistical analysis was conducted using GraphPad Prism. Details of biological replicates and statistical tests are provided in the corresponding figure legends. All data sets were tested for normality using the Shapiro-Wilk test and were confirmed to meet the normality criteria. Statistical analyses were performed as appropriate based on the experimental design.

## Acknowledgements

This work was funded by NIH (1U19AI158080, MM, MK). We thank David Weiss, Dan Andersson, Bruce Levin and Daniel Weissman for helpful discussions throughout the projects.

## Author Contributions

MM and MK conceived the study. MM designed and carried out the experiments and analyzed the data. MK secured funding and provided resources. MM and MK wrote the manuscript. All authors read and approved the manuscript.

## Data Availability Statements

Source data for figures are provided in the Source Data file.

