## Supplementary Material for "Cryptic Evolution of Heteroresistance as Adaptation to Treatment Interruptions"

**Supplementary Text.**

**Derivation of the Recovery-Time Condition Favoring heteroresistance (HR) over monoresistance (MR)**

We consider an environmental cycle consisting of an antibiotic phase of duration $T_{on}$, followed by an antibiotic-free recovery phase $T_{recovery}$. Let $N_{\mathrm{MR}}$ and $N_{\mathrm{HR}}$ denote the total population sizes of the MR and HR strains at the beginning of the antibiotic phase, respectively.

**Survival and growth during antibiotic exposure**

During the antibiotic phase, only resistant cells survive. Let $RF_{\mathrm{MR}}$ and $RF_{\mathrm{HR}}$ denote the resistant-cell survival fractions of MR and HR, respectively. For MR, $RF_{\mathrm{MR}} \approx 1$, whereas it is much lower for HR (e.g., $RF_{\mathrm{HR}} \approx 10^{-3}$–$10^{-4}$). Our analysis of growth rates in the presence of antibiotic shows that the population growth rate of MR is similar to the growth rate of a resistant subpopulation in HR (Fig. 3a and 3c). We therefore define a common growth rate of resistant cells, $\lambda_{on}$.

After time $T_{on}$ under antibiotic treatment, the surviving populations are therefore

$$N'_{\mathrm{MR}} = N_{\mathrm{MR}}\, RF_{\mathrm{MR}}\, e^{\lambda_{on} T_{on}}, \qquad N'_{\mathrm{HR}} = N_{\mathrm{HR}}\, RF_{\mathrm{HR}}\, e^{\lambda_{on} T_{on}}. \qquad \text{Eq. S1}$$

**Growth during the antibiotic-free phase**

In the antibiotic-free environment, the two strains grow with distinct rates: $\lambda_{\mathrm{MR},0}$ and $\lambda_{\mathrm{HR},0}$ (Fig. 3a).

After growth for duration $T_{recovery}$, the populations become

$$N''_{\mathrm{MR}} = N'_{\mathrm{MR}} e^{\lambda_{\mathrm{MR},0} T_{recovery}} = N_{\mathrm{MR}}\, RF_{\mathrm{MR}}\, e^{\lambda_{on} T_{on} + \lambda_{\mathrm{MR},0} T_{recovery}}, \qquad \text{Eq. S2}$$

and

$$N''_{\mathrm{HR}} = N'_{\mathrm{HR}} e^{\lambda_{\mathrm{HR},0} T_{recovery}} = N_{\mathrm{HR}}\, RF_{\mathrm{HR}}\, e^{\lambda_{on} T_{on} + \lambda_{\mathrm{HR},0} T_{recovery}}. \qquad \text{Eq. S3}$$

The factors

$$G_{\mathrm{MR}} = RF_{\mathrm{MR}}\, e^{\lambda_{on} T_{on} + \lambda_{\mathrm{MR},0} T_{recovery}}, \qquad \text{Eq. S4}$$

and

$$G_{\mathrm{HR}} = RF_{\mathrm{HR}}\, e^{\lambda_{on} T_{on} + \lambda_{\mathrm{HR},0} T_{recovery}}\ . \qquad \text{Eq. S5}$$

represent the per-cycle multiplicative growth of the two strains.

**Condition for HR to increase relative to MR**

HR has a selective advantage over MR if its per-cycle growth factor exceeds that of MR, i.e., $G_{\mathrm{HR}} > G_{\mathrm{MR}}$. Substituting the expressions above and canceling the common term $e^{\lambda_{on} T_{on}}$ yields $RF_{\mathrm{HR}}\, e^{\lambda_{\mathrm{HR},0} T_{recovery}} > RF_{\mathrm{MR}}\, e^{\lambda_{\mathrm{MR},0} T_{recovery}}$. Rearranging it, we have

$$(\lambda_{\mathrm{HR},0} - \lambda_{\mathrm{MR},0}) T_{recovery} > \ln\left(\frac{RF_{\mathrm{MR}}}{RF_{\mathrm{HR}}}\right). \qquad \text{Eq. S6}$$

Thus, HR increases from cycle to cycle when the antibiotic-free period exceeds the threshold

$$T_{recovery} > \frac{\ln\left(RF_{\mathrm{MR}}/RF_{\mathrm{HR}}\right)}{\lambda_{\mathrm{HR},0} - \lambda_{\mathrm{MR},0}}. \qquad \text{Eq. S7}$$

By definision, $RF_{\mathrm{MR}} \approx 1$.

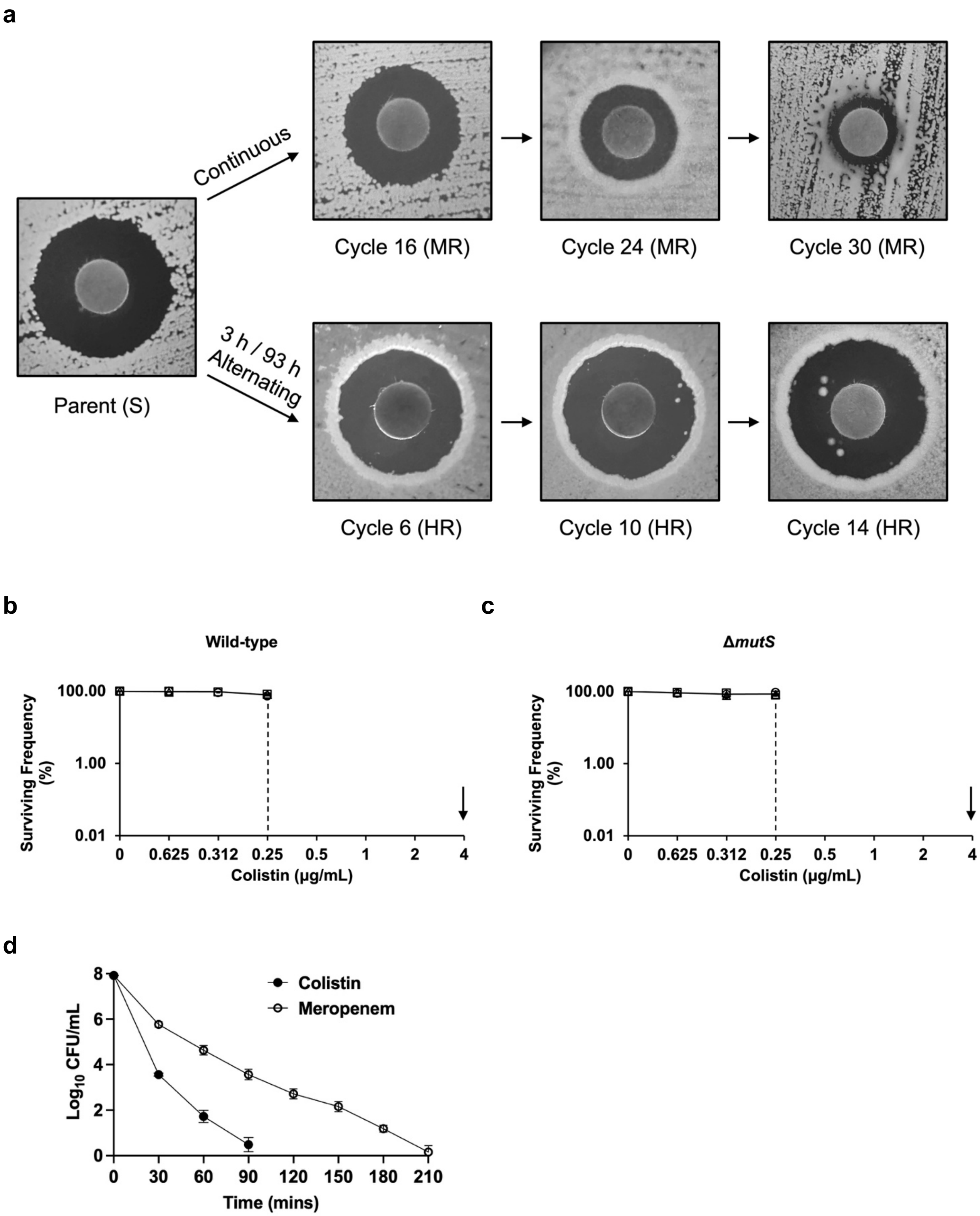

a
Continuous
3 h / 93 h
Alternating
Parent (S)
Cycle 16 (MR)
Cycle 24 (MR)
Cycle 30 (MR)
Cycle 6 (HR)
Cycle 10 (HR)
Cycle 14 (HR)
b
Wild-type
Surviving Frequency (%)
100.00
1.00
0.01
0
0.625
0.312
0.25
0.5
1
2
4
Colistin (μg/mL)
c
ΔmutS
Surviving Frequency (%)
100.00
1.00
0.01
0
0.625
0.312
0.25
0.5
1
2
4
Colistin (μg/mL)
d
Colistin
Meropenem
Log10 CFU/mL
8
6
4
2
0
0
30
60
90
120
150
180
210
Time (mins)

**Supplementary Fig. 1. Colistin susceptibility of parental and evolved strains.**

(a) Disk diffusion assays were conducted as described in Methods using custom-made disks containing 64 µg colistin. The parental strain (Cycle 0) is shown alongside representative strains evolved under continuous exposure (Cycles 16, 24, and 30; top panel) and alternating exposure ($T_{on}/T_{off}$ = 3 / 93 hours, Cycles 6, 10, and 14; bottom panel).

(b,c) Population analysis profiling (PAP) curves for *E. coli* K-12 NCM3722 and its Δ*mutS* derivative. Deletion of *mutS*, used to accelerate evolution, does not alter baseline colistin susceptibility, as both strains show sharp cutoffs well below the clinical breakpoint (4 µg/mL, arrow). Symbols represent independent cultures (n = 3); lines and error bars indicate mean ± SD.

(d) Time–kill curves of the parental strain exposed to colistin or meropenem at their clinical breakpoint concentrations (4 µg/mL). Susceptible cells were reduced to near-complete clearance within ~1.5 h for colistin and ~3.5 h for meropenem. To ensure reliable elimination of susceptible cells, we conservatively set the minimum $T_{on}$ in the evolution experiments to 3 h for colistin and 6 h for meropenem.

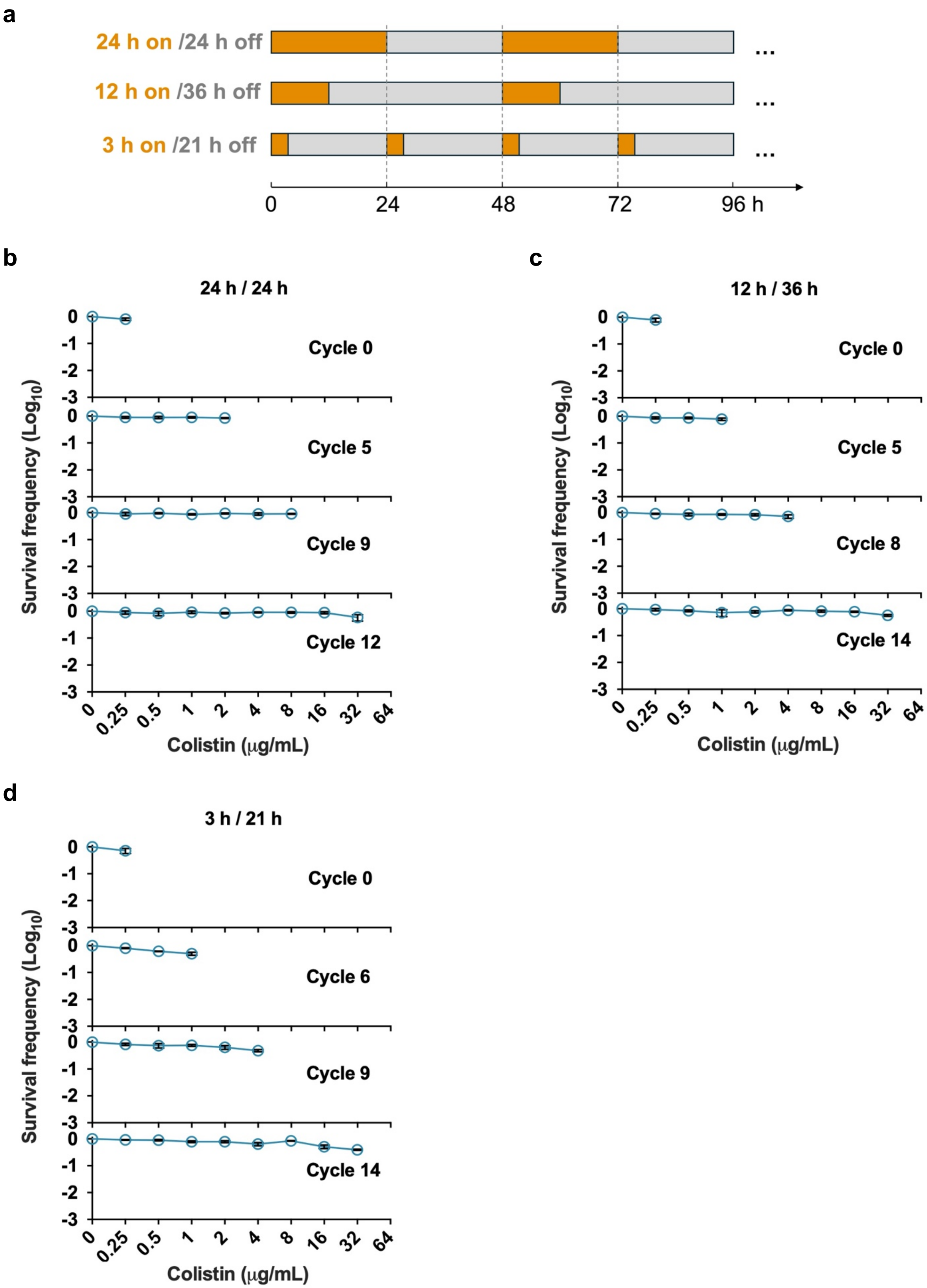

a
24 h on /24 h off
12 h on /36 h off
3 h on /21 h off
...
0
24
48
72
96 h
b
24 h / 24 h
Cycle 0
Cycle 5
Cycle 9
Cycle 12
Survival frequency (Log10)
Colistin (μg/mL)
0
0.25
0.5
1
2
4
8
16
32
64
c
12 h / 36 h
Cycle 0
Cycle 5
Cycle 8
Cycle 14
Survival frequency (Log10)
Colistin (μg/mL)
d
3 h / 21 h
Cycle 0
Cycle 6
Cycle 9
Cycle 14
Survival frequency (Log10)
Colistin (μg/mL)

**Supplementary Fig. 2. Evolution of mono-resistance under short-cycle exposure.**

a) *E. coli* was subjected to frequent alternating cycles of colistin exposure and drug-free phases. b-d) PAP curves of cells evolved under 24 h colistin / 24 h colistin-free phases (b), 12 h / 36 h (c), and 3 h / 21 h (d). Sharp cut-offs in the PAP curves indicate the evolution of mono-resistance. Different symbols represent independent replicate cultures. Black lines and error bars indicate the mean and standard deviation across three replicates.

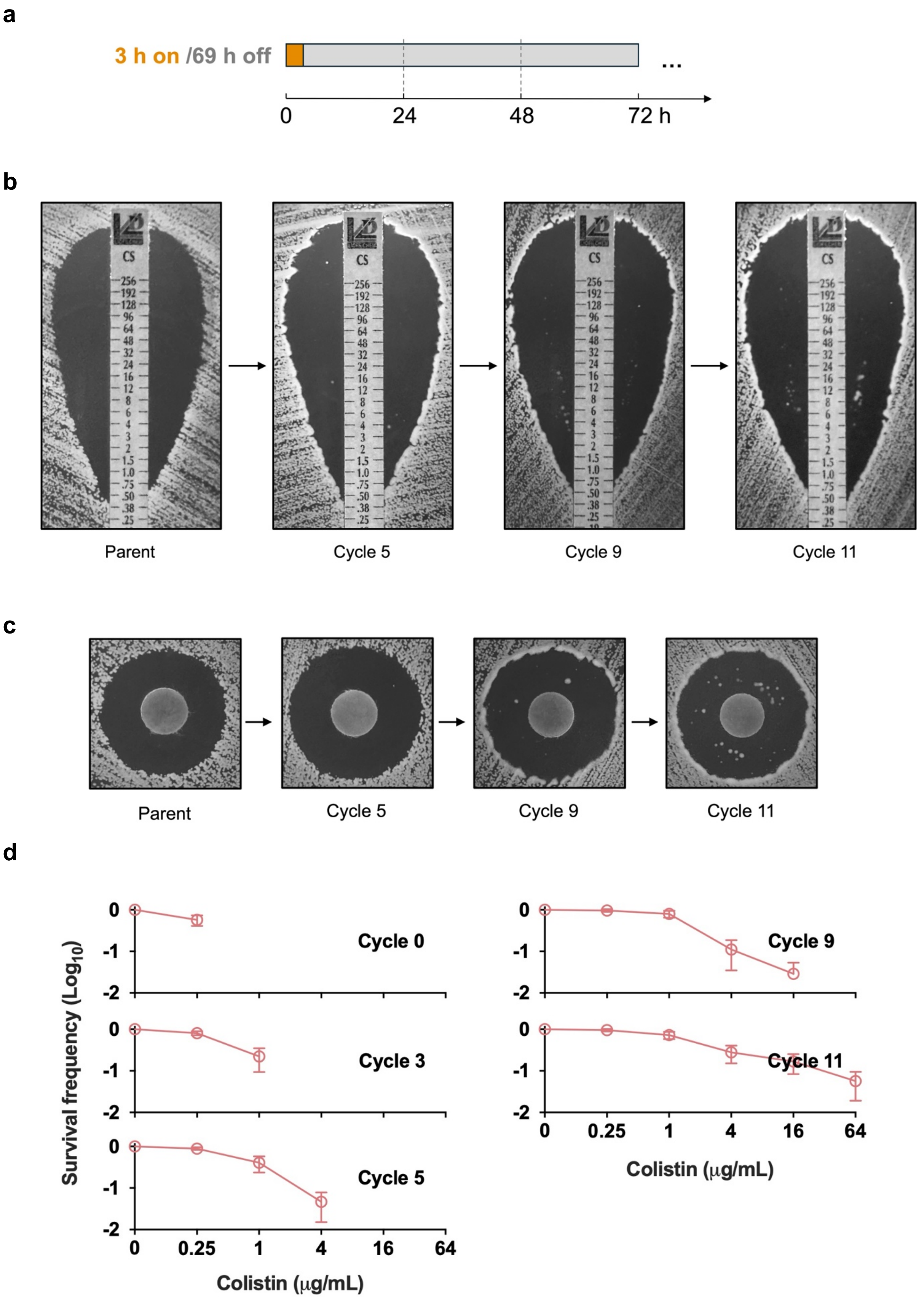
a
3 h on /69 h off
...
0
24
48
72 h
b
CS
Parent
Cycle 5
Cycle 9
Cycle 11
c
Parent
Cycle 5
Cycle 9
Cycle 11
d
Survival frequency (Log10)
Cycle 0
Cycle 3
Cycle 5
Cycle 9
Cycle 11
0
-1
-2
0
0.25
1
4
16
64
Colistin (μg/mL)

**Supplementary Fig. 3. Evolution of HR under 3 h / 69 h alternating exposure.**

a) *E. coli* was subjected to 3 h / 69 h alternating cycles of colistin exposure and drug-free phases.

b) Representative inhibition-zone images of the parental strain and isolates evolved under 3 h / 69 h alternating regimen.

c) Disk diffusion assays were performed as described in Methods using custom-made disks containing 64 μg colistin. The parental strain (Cycle 0) is shown alongside representative evolved strains evolved from Cycles 5, 9, and 11 ($T_{on}/T_{off}$ = 3 h / 69 h).

d) PAP curves of isolates evolved under 3 h colistin / 69 h colistin-free exposure. Symbols and error bars indicate the mean and standard deviation from five independent replicates.

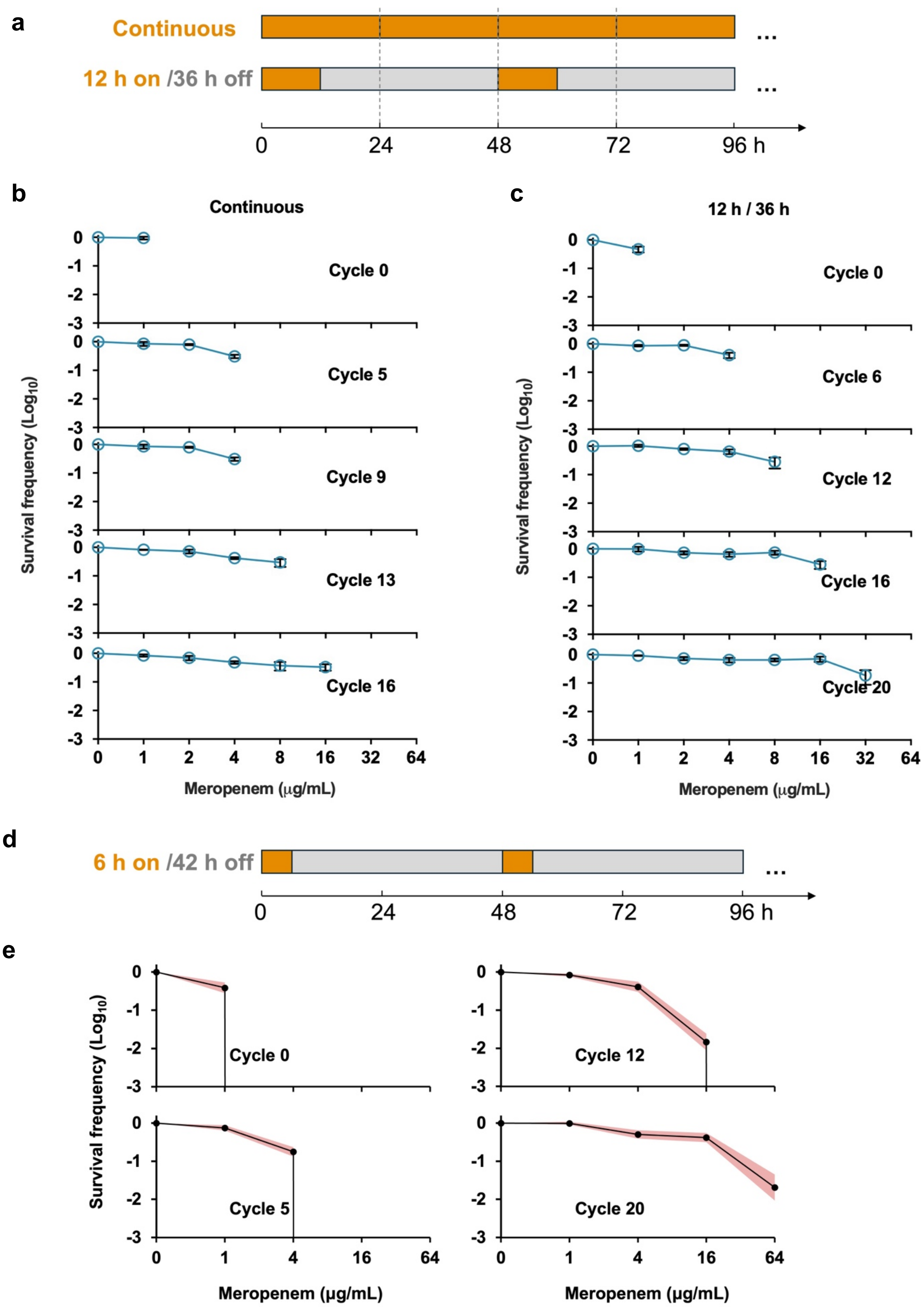
a
Continuous
12 h on /36 h off
0
24
48
72
96 h
b
Continuous
c
12 h / 36 h
Cycle 0
Cycle 5
Cycle 9
Cycle 13
Cycle 16
Cycle 0
Cycle 6
Cycle 12
Cycle 16
Cycle 20
Survival frequency (Log10)
Meropenem (μg/mL)
d
6 h on /42 h off
e
Cycle 0
Cycle 5
Cycle 12
Cycle 20

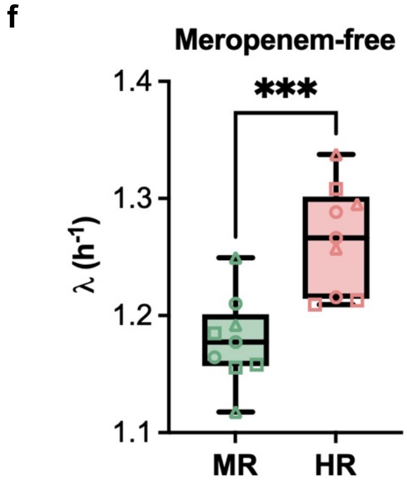


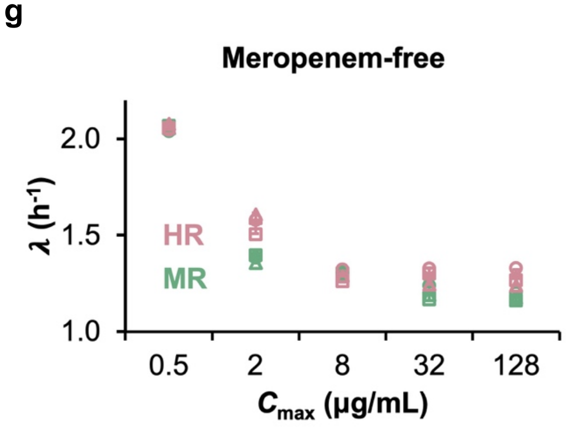


**Supplementary Fig. 4. Evolution under meropenem treatment.**

a-c) *E. coli* was subjected to either continuous meropenem exposure or alternating cycles of meropenem (12hr) and drug-free phases (36 hr). PAP curves reveal the evolution of mono-resistance. Different symbols represent independent replicate cultures. Black lines and error bars indicate the mean and standard deviation across three replicates.

d,e) *E. coli* was subjected to alternating cycles of meropenem (6 hours) and extended drug-free phases (42 hours). Heteroresistance (HR) emerged, as evidenced by the gradual decline in survival frequency in the PAP curves. According to a standard criterium [1,2,3], isolates at Cycle 12 and 20 are classified as HR. Solid dots represent the mean survival frequency of independent colonies (n=50) from five parallel cultures, demonstrating that HR evolution is reproducible; shaded regions indicate the full range.

f) Growth rates ($\lambda$) of meropenem-MR and HR strains from Cycle 16 and Cycle 13 respectively. Three biological replicates were tested per strain. ***$p < 0.001$, based on two-way ANOVA with multiple comparisons.

g) We measured the antibiotic-free growth rates of HR and MR strains throughout the evolution experiment. Resistance level was quantified by the minimum inhibitory concentration (MIC), defined as the lowest colistin concentration that completely inhibited population growth. Growth rates measured in drug-free conditions were then plotted against MIC.

**a**

**Cycle 0**
**(MIC = 0.5 μg/mL)**

| Postion | Mutation | Annotation | Description |
|---|---|---|---|
| 785,524 | A→C | intergenic | phosphoribosylaminoimidazolecarboxamide formyltransferase/16S ribosomal RNA |
| 1,496,971 | A→T | L110Q | transposase |
| 1,974,562 | G→A | V18M | hypothetical protein |
| 2,030,437 | $(A)_{5\rightarrow 4}$ | intergenic | hypothetical protein |
| 2,350,314 | T→C | Y20H | hypothetical protein |
| 2,679,979 | A→G | L296L | transposase |
| 3,544,701 | C→T | intergenic | ecotin/malare:quinone oxidoreductase |
| 3,971,655 | G→A | noncoding | 16S ribosomal RNA |

**Cycle 3**
**(MIC = 2 μg/mL)**

| Postion | Mutation | Annotation | Description |
|---|---|---|---|
| 785,528 | Δ1bp | intergenic | phosphoribosylaminoimidazolecarboxamide formyltransferase/16S ribosomal RNA |
| 1,903,290 | C→A | intergenic | asparagine synthetase B/UMP phosphatase |
| 1,911,762 | A→G | intergenic | glutamate--tRNA ligase/chitoporin |
| 1,941,943 | C→T | intergenic | H repeat-associated protein YdcC/YbfD |

**Cycle 6**
**(MIC = 8 μg/mL)**

| Postion | Mutation | Annotation | Description |
|---|---|---|---|
| 1,940,870 | T→C | F26F | H repeat-associated protein YdcC |
| 3,808,980 | C→T | intergenic | ethanolamine utilization protein EutH/EutG |

**Cycle 10**
**(MIC = 32 μg/mL)**

| Postion | Mutation | Annotation | Description |
|---|---|---|---|
| 910,452 | G→A | C84Y | sensor protein PmrB |
| 1,325,672 | Δ6bp | ΔEY199-200 | UDP-3-O-[3-hydroxymyristoyl] N-acetylglucosamine deacetylase LpxC |

**Cycle 14**
**(MIC = 128 μg/mL)**

**b**

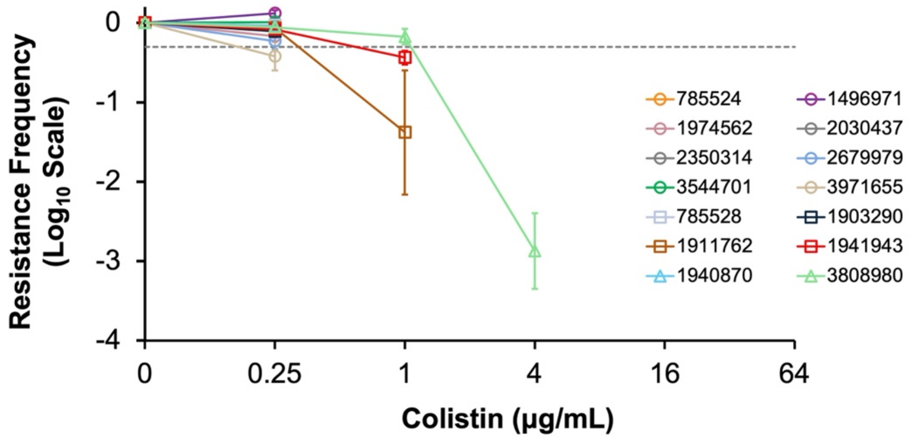

**Supplementary Fig. 5. Mutations identified in heteroresistant strains.**

(a) Whole-genome sequencing of strains evolved under 3 h on/93 h off colistin cycles was conducted. The table summarizes mutation positions, nucleotide changes, gene annotations, and predicted functions.

(b) The corresponding mutations were introduced into the genome of wild-type *E.coli* via CRISPR, and PAP assays were performed to assess their impact on colistin resistance. Data are shown as mean ± s.d., n=3.

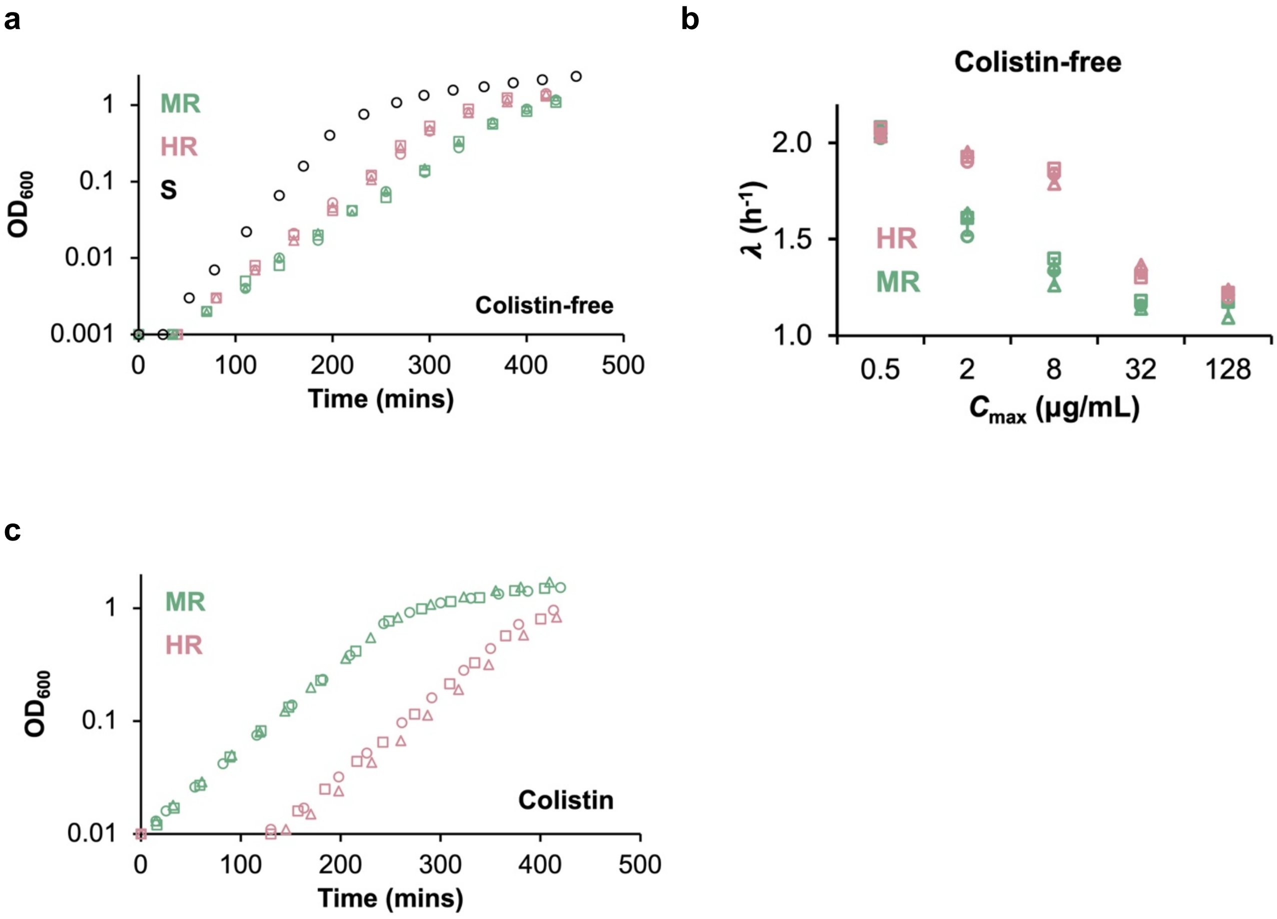


**Supplementary Fig. 6. Growth curves of heteroresistant (HR) and mono-resistant (MR) strains.**

a, c) Growth curves of MR and HR strains from Cycle 23 and 10 respectively, measured under colistin-free condition (a) and in 16 μg/mL colistin (c). Different symbols represent independent replicate cultures (n=3). Dashed lines fitting with the growth curves indicate exponential phase.

b) We measured the antibiotic-free growth rates of HR and MR strains throughout the evolution experiment. Resistance level was quantified by the minimum inhibitory concentration (MIC), defined as the lowest colistin concentration that completely inhibited population growth. Growth rates measured in drug-free conditions were then plotted against MIC.

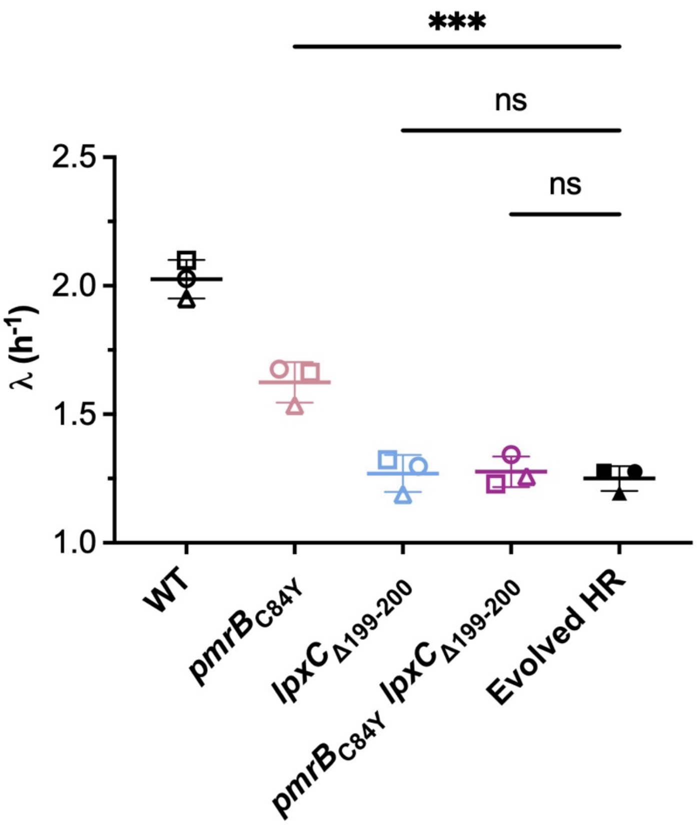


**Supplementary Fig. 7. *pmrB*$_{\mathbf{C84Y}}$ and *lpxC*$_{\mathbf{\Delta EY199–200}}$ mutations recapitulate the fitness cost of the HR strain under antibiotic-free conditions.**

Growth rates ($\lambda$, $h^{-1}$) of wild-type (WT), *pmrB*$_{C84Y}$, *lpxC*$_{\Delta EY199–200}$, the corresponding double mutant, and the evolved colistin-HR strain in Cycle 14 were measured under antibiotic-free conditions. Both mutants exhibited reduced growth rates relative to WT, comparable to the fitness cost observed in the evolved HR strain. Data are shown as mean ± s.d., n=3.

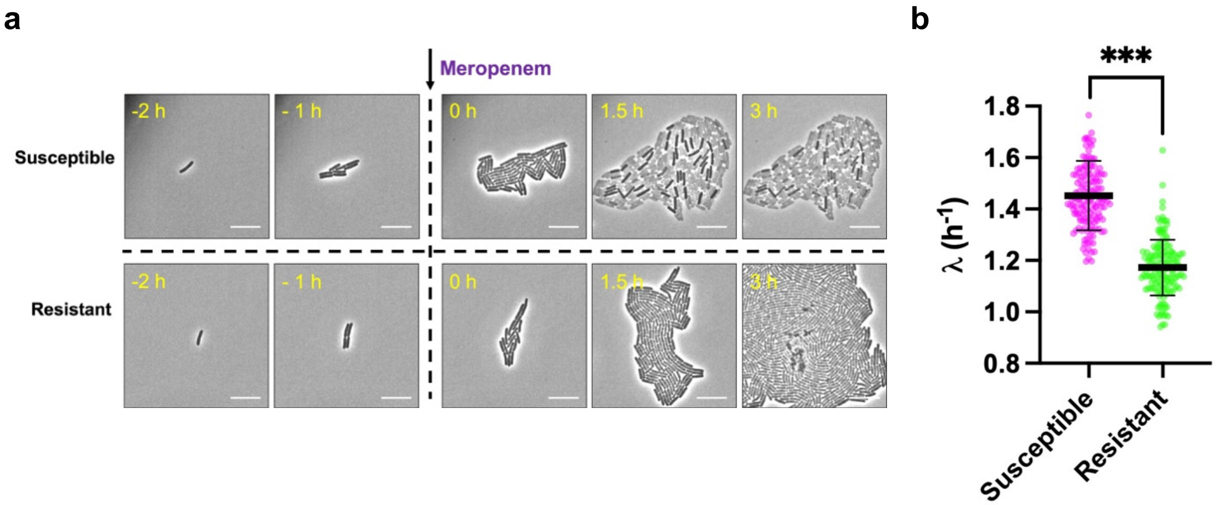


**Supplementary Fig. 8. Fitness analysis of lab-evolved meropenem hetero-resistant (HR) strain at single cell level.**

(a) Images to the left of the arrow show the pre-treatment phase; those to the right show the post-treatment phase. Cells were classified based on their response to the treatment: susceptible (killed at 2 µg/mL) or resistant (surviving at 16 µg/mL). Scale bar: 10 µm.

(b) Susceptible cells exhibited significantly higher growth rates prior to meropenem treatment. Three independent experiments were conducted, with 50 cells per group per replicate. Black lines and error bars indicate the mean and standard deviation. ***$p < 0.001$, based on unpaired t-test.

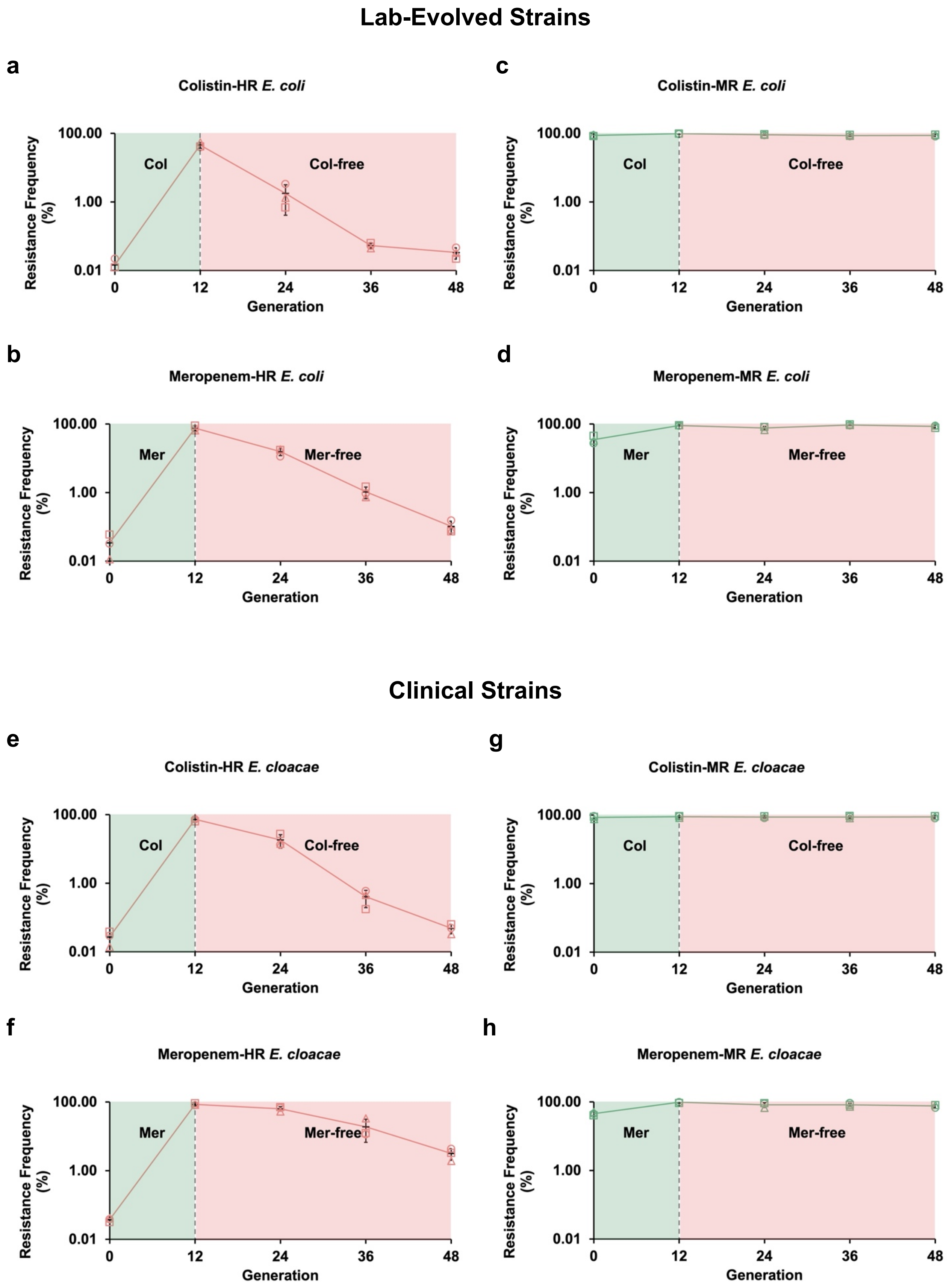
Lab-Evolved Strains
a
Colistin-HR E. coli
Col
Col-free
c
Colistin-MR E. coli
b
Meropenem-HR E. coli
Mer
Mer-free
d
Meropenem-MR E. coli
Clinical Strains
e
Colistin-HR E. cloacae
g
Colistin-MR E. cloacae
f
Meropenem-HR E. cloacae
h
Meropenem-MR E. cloacae
Resistance Frequency (%)
100.00
1.00
0.01
Generation
0
12
24
36
48

**Supplementary Fig. 9. Dynamic population restructuring of hetero-resistant (HR) strains**

**(a, b)** Population analysis profiling (PAP) of lab-evolved *E. coli* strains exhibiting colistin-HR or meropenem-HR phenotypes (MIC = 32 µg/mL for either drug). PAP assays were performed at 16 µg/mL to measure the frequency of highly resistant subpopulations. The basal frequency ranged from 0.01% to 0.1%. Upon culturing in 16 µg/mL of colistin or meropenem (green region), the frequency increased to nearly 100%. When the cultures were transferred to drug-free LB medium (pink region), the resistance frequency reverted to basal levels. Different symbols represent independent replicate cultures; black lines and error bars indicate the mean and standard deviation.

**(c, d)** Equivalent experiments performed with lab-evolved colistin-MR or meropenem-MR *E. coli* strains.

**(e–h)** The experimental protocol described above for lab-evolved *E. coli* strains was identically applied to clinical HR and MR *E. cloacae* strains.

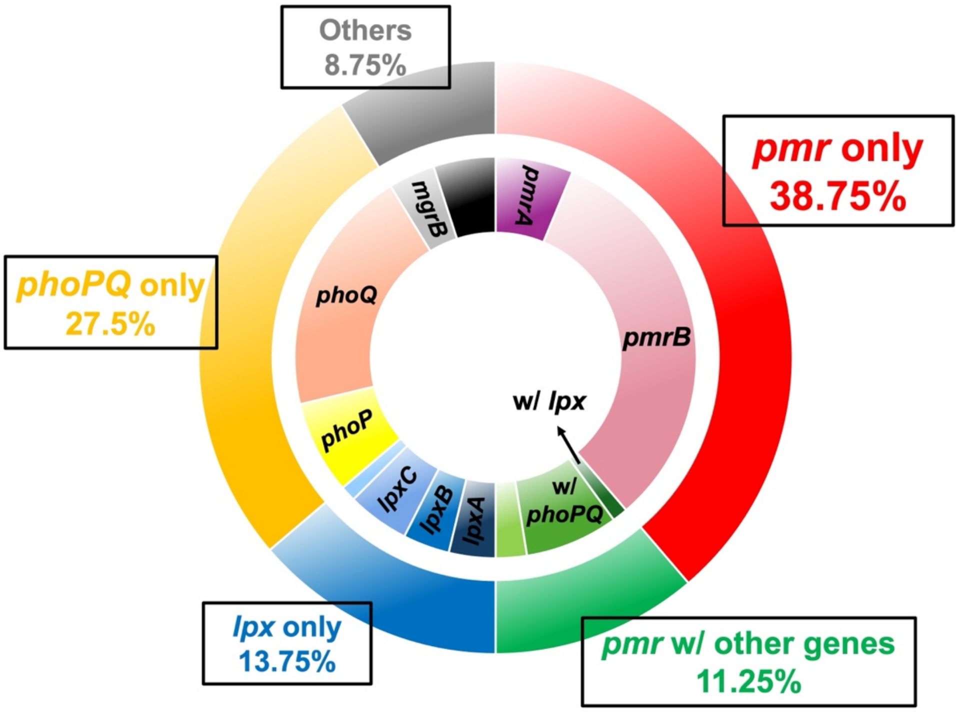


**Supplementary Fig. 10. Frequency of mutations found in previous studies of colistin-heteroresistant clinical isolates.**

We surveyed prior reports of colistin heteroresistance in clinical isolates and underlying mutations (Supplementary Table 1). Here, we report the percentage distribution of major mutations. The outer ring represents operon-level categories of mutations, including *pmr* only, *pmr* with other genes, *lpx* only, *phoPQ* only, and other genes only. The inner ring denotes finer subcategories within each major class. Specifically, the *pmr* only category comprises mutations in *pmrA* and *pmrB*; *pmr* with other genes includes co-occurring mutations with *lpxC*, *phoPQ*, *mgrB*, or other genes; *lpx* only includes mutations in *lpxA*, *lpxC*, *lpxD*, and *lpxM*; *phoPQ* only includes mutations in *phoP* and *phoQ*; Others include mutations in genes such as *mgrB* and *crrS*, among others.

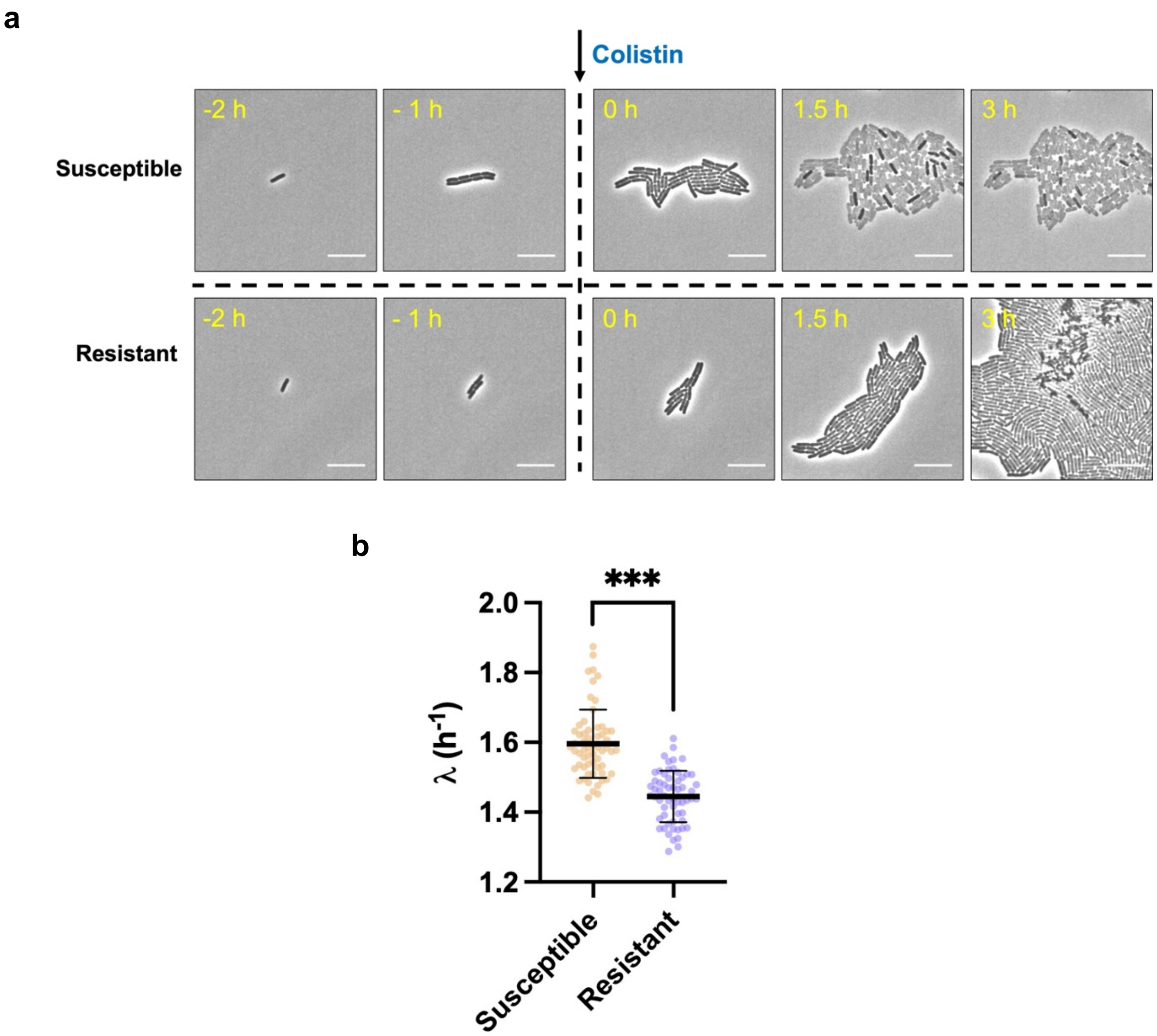


**Supplementary Fig. 11. Fitness analysis of clinical colistin hetero-resistant (HR) *E. cloacae* strain at single cell level.**

(a) Images to the left of the arrow show the pre-treatment phase; those to the right show the post-treatment phase. Cells were classified based on their response to the treatment: susceptible (killed at 16 μg/mL) or resistant (surviving at 16 μg/mL). Scale bar: 10 μm.

(b) Susceptible cells exhibited significantly higher growth rates prior to colistin treatment. Three independent experiments were conducted, with 20 cells per group per replicate. Black lines and error bars indicate the mean and standard deviation. ***$p < 0.001$, based on unpaired t-test.

### Supplementary Table 1. Clinical isolates tested in this study[13, 14].

| Strain ID | Genus | Species | Colistin Susceptibility | Strain ID | Genus | Species | Colistin Susceptibility |
|---|---|---|---|---|---|---|---|
| AMK107 | *Enterobacter* | *cloacae* | HR | KMK143 | *Enterobacter* | *aerogenes* | S |
| KMK100 | *Enterobacter* | *cloacae* | HR | KMK144 | *Enterobacter* | *aerogenes* | S |
| KMK101 | *Enterobacter* | *asburiae/ludwigii* | HR | KMK145 | *Enterobacter* | *aerogenes* | S |
| KMK102 | *Enterobacter* | *asburiae* | HR | KMK146 | *Enterobacter* | *cloacae* | S |
| KMK103 | *Enterobacter* | *ludwigii* | HR | KMK147 | *Enterobacter* | *aerogenes* | S |
| KMK104 | *Enterobacter* | *kobei* | HR | KMK148 | *Enterobacter* | *cloacae* | S |
| KMK105 | *Enterobacter* | *cloacae* | HR | KMK149 | *Enterobacter* | *cloacae* | S |
| KMK106 | *Enterobacter* | *cloacae/kobei* | HR | KMK150 | *Enterobacter* | *aerogenes* | S |
| KMK107 | *Enterobacter* | *cloacae* | HR | KMK151 | *Enterobacter* | *cloacae* | S |
| KMK110 | *Enterobacter* | *cloacae/asburiae* | HR | KMK152 | *Enterobacter* | *cloacae* | S |
| KMK111 | *Enterobacter* | *cloacae* | HR | KMK153 | *Enterobacter* | *cloacae* | S |
| KMK114 | *Enterobacter* | *aerogens* | HR | KMK154 | *Enterobacter* | *aerogenes* | S |
| KMK115 | *Enterobacter* | *cloacae/asburiae* | HR | KMK155 | *Enterobacter* | *cloacae* | S |
| KMK116 | *Enterobacter* | *kobei* | HR | KMK156 | *Enterobacter* | *cloacae* | S |
| KMK121 | *Enterobacter* | *cloacae* | HR | KMK157 | *Enterobacter* | *cloacae* | S |
| KMK124 | *Enterobacter* | *cloacae/asburiae* | HR | KMK158 | *Enterobacter* | *cloacae* | S |
| KMK127 | *Escherichia* | *coli* | HR | KMK159 | *Enterobacter* | *hormaechei/cloacae* | S |
| KMK128 | *Escherichia* | *coli* | HR | KMK160 | *Enterobacter* | *cloacae* | S |
| KMK129 | *Escherichia* | *coli* | HR | KMK161 | *Enterobacter* | *aerogenes* | S |
| AMK104 | *Enterobacter* | *cloacae* | R | KMK162 | *Enterobacter* | *cloacae/hormaechei* | S |
| KMK130 | *Enterobacter* | *cloacae* | R | KMK163 | *Enterobacter* | *cloacae* | S |
| KMK131 | *Enterobacter* | *kobei/asburiae* | R | KMK164 | *Enterobacter* | *cloacae* | S |
| KMK132 | *Enterobacter* | *aerogenes* | R | KMK165 | *Enterobacter* | *cloacae* | S |
| KMK133 | *Enterobacter* | *cloacae/asburiae* | R | KMK166 | *Enterobacter* | *cloacae* | S |
| AMK105 | *Enterobacter* | *cloacae* | S | KMK167 | *Enterobacter* | *cloacae* | S |
| KMK135 | *Enterobacter* | *cloacae* | S | KMK168 | *Enterobacter* | *cloacae* | S |
| KMK136 | *Enterobacter* | *aerogenes* | S | AMK105 | *Enterobacter* | *cloacae* | S |
| KMK137 | *Enterobacter* | *aerogenes* | S | KMK170 | *Escherichia* | *coli* | S |
| KMK138 | *Enterobacter* | *aerogenes* | S | KMK171 | *Escherichia* | *coli* | S |
| KMK139 | *Enterobacter* | *cloacae* | S | KMK172 | *Escherichia* | *coli* | S |
| KMK140 | *Enterobacter* | *cloacae* | S | KMK83 | *Klebsiella* | *pneumoniae* | HR |
| KMK141 | *Enterobacter* | *cloacae* | S | KMK89 | *Klebsiella* | *pneumoniae* | HR |
| KMK142 | *Enterobacter* | *cloacae* | S | KMK95 | *Klebsiella* | *pneumoniae* | HR |

### Supplementary Table 2. Mutations demonstrated to confer colistin heteroresistance in previous studies.

| Species | PmrA | PmrB | PmrC | LpxA | LpxC | LpxD | LpxM | PhoP | PhoQ | MgrB | YciM | MutS | CrrA | ParR | CprS | References |
|---|---|---|---|---|---|---|---|---|---|---|---|---|---|---|---|---|
| *E. coli* | | C84Y | | | ΔE199-Y200 | | | | | | | | | | | This study |
| | | R93P | | | | | | | | | | | | | | Kuang et al., 2020 [1] |
| | | L14R | | | | | | | | | | | | | | Liao et al., 2020 [2] |
| | | P95L | | | | | | | | | | | | | | |
| *P. aeruginosa* | | V28A | | | | | | | | | | | | | | Kapel et al., 2022 [3] |
| | | P254S | | | | | | | | | | | | | | |
| | | V199I,S257N | | | | | | | | | | | | R146H | V181I,R209L | Lin et al., 2019 [4] |
| | | V185A | | | | | | | | | | | | | | |
| | | V15I,G68S | | | | | | | | | | | | | | |
| | | G179D,I349V | | | | | | | | | | | | | | |
| | | | | | | | | | V260G | | | | | | | |
| | | G179D | | | | | | | V260G | | | | | | | |
| | | D45E | | | | | | | | | | | | | | |
| | | A190G | | | | | | | | | | | | | | |
| | | S27R | | | | | | | | | | | | | | |
| *A. baumannii* | | M308R | | | | | | | | | | | | | | Charretier et al., 2018 [5] |
| | | S144KLAGS | | | | | | | | | | | | | | |
| | M12I | | | | | | | | | | | | | | | |
| | | P170L | | | | | | | | | | | | | | |
| | A14V | | | | | | | | | | | | | | | Hong et al., 2020 [6] |
| | | E301D | | | | | | | | | | | | | | |
| | | | | M169T | | | | | | | | | | | | |
| | | | | R258H | | | | | | | | | | | | |
| | | | | T81M | | | | | | | | | | | | |
| | | | | | | T290I | | | | | | | | | | |
| | | | | | | G231V | | | | | | | | | | |
| | | T235I | | | | | | | | | | | | | | |
| | | L271F | | | | | | | | | | | | | | |
| | | P170L | | | | | | | | | | | | | | |
| | | | | | M62I | | | | | | | | | | | |
| | | L271F | | | | | | | | | | | | | | |
| | | S17R | | | | | | | | | | | | | | |
| | | | | | C63Y | | | | | | | | | | | |
| | | G260D | | | | | | | | | | | | | | |
| | | | | | | G309V | | | | | | | | | | |
| | S16L | | | | | | | | | | | | | | | |
| | | T232I | | | | | | | | | | | | | | |
| | | L168S | | | | | | | | | | | | | | |
| | | | | | | G275* | | | | | | | | | | Hematian et al., 2025 [7] |
| | | A138T | | | | | | | | | | | | | | |
| | | E301D | | | | T289I | | | | | | | | | | |
| | | | | | P46Q | | | | | | | | | | | |
| *E. cloacae* | | | | | | | | | R423C | | | | | | | Wang et al., 2025 [8] |
| | | | | | | | | | L9FS | | | | | | | |
| | | | | | | | | | N90D,N150D | | | | | | | |
| | | | | | | | | | N90D | | | | | | | |
| | | | | | | | | | G459C | | | | | | | Wei et al., 2025 [9] |
| | | | | | | | | | L250Q | | | | | | | |
| | | | | | | | | | Q20K | | | | | | | |
| | | | | | | | | | C392Y | | | | | | | |
| | | | | | | | | | G385E | | | | | | | |
| | | | | | | | | | R294H | | | | | | | |
| *K. pneumoniae* | | | | | | | | D191Y | | | | | | | | Jayol et al., 2015 [10] |
| | | | | | | | | | | | V43G | | | | | |
| | | | | | | | | | A21S | | | | | | | |
| | | | | | | | V30G | | | | | | | | | |
| | | | | | | | | | | 38G ins | | | | | | Bardet et al., 2017 [11] |
| | R203K | D150N | | | | | | R198H,K199N | L414R | | | | | | | Cheong et al., 2019 [12] |
| | R203K | D150N | | | | | | | D152N | | | | | | | |
| | I178F | | | | | | | | | | | | | | | |
| | | D150N | | | | | | | | | | | | | | |
| | R203K | | | | | | | | | | | | | | | |
| | R203K | | | | | | | | L414R | | | | | | | |
| | | | | | | | | | | | | K103* | | | | Sato et al., 2020 [13] |
| | | | | | | | | | A351D | | | | | | | Morales-León et al., 2020 [14] |
| | | | | | | | | T104A | | | | | | | | |
| | | P95L | | | | | | T104A | | | | | | | | |
| | | R256G | | | | | | | | C39* | | | | | | |
| | | | | | | | | | V24G | | | | | | | Sánchez-León et al., 2023 [15] |
| | | | | | | | | | G385V | | | | | | | |
| | | | | | | | | | | ΔW6 | | | | | | |
| | | | | | | | | | | | | | D96E | | | |
| | | | | | | | | | | | | | D96N | | | |
| | | | F27C, S257L, R319Q | | | | | | A284V | | | | | | | |
| | | Q202K | | | | | | | | | | | | | | |
| | | | | | | | | Y98C,V134Y | | | | | | | | |
| | | | | | | | | | | ΔS36K / ΔG37-W47 | | | | | | |
| | | | | | | | | | G385C | | | | | | | |
| | | | | | | | | Y98C,S128P | | | | | | | | |
| | | | | | | | | Y98C,V126A | | | | | | | | |
| | | | | | | | | Y98C,G166D | | | | | | | | |

**Supplementary Table 3.**

| **Strains Name** | **Description** | **Source** |
|---|---|---|
| NMK1 | *E. coli* NCM3722, "wild-type" | Kim lab collection |
| NMK99 | NMK1 with plasmid pKD46, AmpR | Kim lab collection |
| NMK476 | Δ*mutS*::km_FRT in NMK1 | This study |

| **Plasmids Name** | **Description** | **Source** |
|---|---|---|
| pKD46 | Encodes lambda red recombinase | Kim lab collection |
| pCas9 | Encodes Cas9 nuclease, tracrRNA and crRNA guide | Addgene |
| pCRISPR | Encodes crRNA targeting a specific sequence | Addgene |

| **Primers Name** | **Sequence** |
|---|---|
| CRISPR_PmrB_Spacer | GACGCTATTTATCTGCTATC |
| CRISPR_lpxC_Spacer | ATCGAATATCTGCAGTCCCG |
| PmrB_GtoA_repair | TGATTGTCCCCGGCGTCTTTATGGTCAGCCTGACGCTATTTATCTACT ATCAGACGGTACGCCGCATCACCCGCCCGCTGGCGGAGCTGC |
| lpxC_6bpDel_repair | CGCCAGATCAGCCGTGCGCGTACGTTCGGTTTCATGCGTGATATCCT GCAGTCCCGTGGTTTGTGCCTGGGCGGCAGCTTCGATTGTGCC |
| PmrB_Pcrispr_Fwd | GAGACCAGTCTCGGAAGCTCGACGCTATTTATCTGCTATC |
| PmrB_Pcrispr_Rev | TTTTTAGAGCTATGCTGTTTGATAGCAGATAAATAGCGTC |
| lpxC_Pcrispr_Fwd | GAGACCAGTCTCGGAAGCTCATCGAATATCTGCAGTCCCG |
| lpxC_Pcrispr_Rev | TTTTTAGAGCTATGCTGTTTCGGGACTGCAGATATTCGAT |
| PmrB_Pcrispr_Seq_Fwd | TCTCGGAAGCTCAAAGGTCT |
| PmrB_Pcrispr_Seq_Rev | AGCAGGACGCACTGACCGAA |
| lpxC_Pcrispr_Seq_Fwd | TGCGCGTACGTTCGGTTTCA |
| lpxC_Pcrispr_Seq_Rev | GCACAATCGAAGCTGCCGCC |
